# Mining for disease-associated microbial metabolites in an age-dependent model of multiple sclerosis

**DOI:** 10.1101/2024.05.27.595846

**Authors:** Annie Pu, Naomi M Fettig, Alexandros Polyzois, Gary YC Chao, Ikbel Naouar, Leah S Hohman, Marissa A Fontaine, Michelle Zuo, Kevin Champagne-Jorgensen, Julia Copeland, Donny Chan, Katherine M Davis, Ruoqi Yu, Sarah J Popple, Natalia A Carranza García, David S Guttman, Kathy D McCoy, Valeria Ramaglia, Frank C Schroeder, Jennifer L Gommerman, Lisa C Osborne

## Abstract

Age is a risk factor for the neurological decline and physical disability that characterize progressive multiple sclerosis (MS). The intestinal microbiota and the bioactive compounds it produces can influence aging, immunity, and the central nervous system (CNS). Here, we use an experimental autoimmune encephalomyelitis (EAE) model that mimics features of progressive MS in aged, but not young, mice to address the intersection of age and the microbiota on EAE outcomes. Although the microbiota of SJL/J mice aged under controlled laboratory conditions does not promote an ‘aged’ non-remitting EAE phenotype, young mice harboring heterochronic human fecal microbiota transplants (hFMT) developed a range of EAE phenotypes. Metabolomic profiling of mice colonized with an aged hFMT that promoted non-remitting EAE indicated a severe reduction in circulating levels of the microbiota-derived tryptophan metabolite indole 3-propionic acid (IPA). IPA-supplementation enforced remission in mice colonized with the non-remitting hFMT, demonstrating the utility of this *in vivo* pipeline for discovering metabolites associated with progressive MS-like disease.

**Summary:** The microbiota is a critical determinant of disease susceptibility in mouse models of MS. Here, Pu & Fettig *et al*. demonstrate that disease outcomes (remitting or non-remitting) are microbiota-responsive, and describe an *in vivo* pipeline that can be mined for microbial metabolites with therapeutic potential for progressive MS.

## Background

Aging is a complex process influenced by both host intrinsic and extrinsic environmental factors. Emerging evidence indicates that the gut microbiota, the community of bacteria, fungi, and viruses that symbiotically inhabit the gastrointestinal tract, changes with advanced age (typically defined as 65+ years) (Ghosh et al., 2022a, 2022b). Invertebrate and small animal studies in controlled environments suggest that aging influences microbiome composition (Cabreiro et al., 2013; Cabreiro and Gems, 2013; Guo et al., 2022; Liu et al., 2012; Thevaranjan et al., 2017), but human studies indicate that external factors such as geography, diet, medication and lifestyle (e.g. independent or community living, social interactions, activity levels) are more indicative of how the microbiota changes than chronological age alone (Claesson et al., 2012; Ghosh et al., 2020). Further, common age-associated co-morbidities (frailty, reduced lifespan and diminished healthspan) have been associated with loss of putative health-promoting microbial taxa and increased presence or abundance of pathobionts (Fettig et al., 2024; Ghosh et al., 2020; Jeffery et al., 2016; Larson et al., 2022; Maffei et al., 2017). The restructured microbiota of healthy and unhealthy agers results in changes to the enzymatic pathways available in the gastrointestinal tract, and production of microbiota-derived metabolites (Rampelli et al., 2013; Ruiz-Ruiz et al., 2020; Wilmanski et al., 2021; Wu et al., 2019). Whether these changes are a cause or consequence of age-associated comorbidities remains unclear.

The gut microbiota can influence tissues distant from the gastrointestinal tract, including the central nervous system (CNS). Microbiota-sensitive pathways of gut-to-CNS communication include trafficking of intestinal leukocytes to the CNS (Pröbstel et al., 2020; Rojas et al., 2019), signals derived from enteric nerves (Cox and Weiner, 2018), and production of microbiota-dependent metabolites that modulate systemic immunity or gain access to the CNS (Erny et al., 2015; Rothhammer et al., 2016). Notably, age-related neurodegenerative conditions such as stroke, Alzheimer’s, and Parkinson’s disease are associated with differences in microbiota composition compared to healthy controls (Fang et al., 2020; Ullah et al., 2023).

Although multiple sclerosis (MS) has not typically been considered an age-related neurodegenerative condition, age is one of the most significant risk factors associated with progressive MS, where neurodegeneration rather than autoimmune inflammation is the primary driver of CNS tissue damage (Lassmann, 2019; Tutuncu et al., 2012). Indeed, people receiving an initial diagnosis later in life are more likely to be diagnosed with primary progressive MS (Kantarci et al., 2016) and the most reliable indicator of disease progression (conversion from relapsing remitting MS) is age (Koch et al., 2007; Tutuncu et al., 2012).

A growing body of evidence indicates that the fecal microbiome of people with MS (pwMS) differs considerably from healthy community or household controls (Cantarel et al., 2015; Chen et al., 2016a; Cosorich et al., 2017; Cox et al., 2021; iMSMS Consortium et al., 2022; Jangi et al., 2016; Miyake et al., 2015; Tremlett et al., 2016). Supporting these epidemiological studies, animal models have demonstrated that the microbiota is an essential regulator of susceptibility to experimental autoimmune encephalomyelitis (EAE). EAE is delayed and severity reduced in germ-free (GF) and antibiotic-treated mice with a depleted microbiota, and susceptibility can be restored by administration of a single CD4^+^ T helper 17 (Th17)-promoting commensal microbe (Lee et al., 2011; Ochoa-Repáraz et al., 2009). Further, intestinal microbiota transplant studies have established a causal role for MS-associated microbiota in disease susceptibility. Ileal and fecal microbiota transplants (FMT) from pwMS into GF mice increased the incidence, accelerated onset, and enhanced EAE severity compared to healthy control microbiota transplants (Berer et al., 2017; Cekanaviciute et al., 2017; Cox et al., 2021; Yoon et al., 2025). Notably, the effects of microbiota on EAE susceptibility are associated with neuroinflammatory T cell activation. Despite the connections between age and MS progression (Tutuncu et al., 2012), age-related changes to the microbiome in other neurodegenerative conditions, and the microbiota in EAE susceptibility (Fettig et al., 2024; Pu et al., 2021), the impact of the microbiota on MS progression, past the initial activation of neuroinflammatory T cells, remains unknown.

We have previously established an EAE model that recapitulates several key pathological features of MS progression that is induced by adoptive transfer (A/T) of encephalitogenic Th17 cells into aged SJL/J mice (Zuo et al., 2022). Compared to young mice that develop a single wave of EAE-induced paralysis followed by remission, aged SJL/J EAE mice exhibit protracted, non-remitting paralysis that is accompanied by enhanced CNS immune infiltration and formation of meningeal tertiary lymphoid tissues (TLT), demyelination, microgliosis, and the emergence of cortical grey matter atrophy (Zuo et al., 2022). This model allows us to separate initial priming of encephalitogenic T cells (i.e., EAE susceptibility) from host-intrinsic factors, like the microbiome, that influence how pre-primed T cells interact with the inflamed CNS to modify disease outcomes – i.e., progression.

We demonstrate herein that chronological aging is not a primary driver of microbiome divergence in SJL/J mice. However, by taking advantage of age-associated differences in the microbiome of healthy human donor pairs, we identified microbial communities that promote a severe, non-remitting “aged EAE” phenotype in young mice. Metabolomic analysis of a candidate aged donor sample that elicited non-remitting EAE in young hosts indicated disrupted tryptophan catabolism, with a striking reduction in indole-3-propionic acid (IPA) production. Exogenous administration of IPA enforced remission in mice colonized with the aged FMT that otherwise drove a progressive MS-like disease, and this was associated with impaired recruitment of “second wave” CD4^+^ Th17 cells to the CNS. Together, these data support a functional role for the microbiota in driving accumulated physical dysfunction in progressive MS-like disease and demonstrate the utility of this *in vivo* SJL/J EAE pipeline to discover metabolites that promote, or could be used to halt, MS progression.

## Results

### Host microbiome dictates the severity and chronicity of Th17 cell transfer EAE

The historic lack of suitable mouse models of grey matter pathology has confounded determination of a role for the microbiota in the pathophysiology of this process. Moreover, whether the host microbiome primarily impacts immune priming or effector activity in the CNS is not distinguished by active immunization models of EAE. We took advantage of observations that adoptive transfer of PLP-primed, Th17 polarized encephalitogenic T cells reliably induces meningeal inflammation and grey matter damage in the brains of naïve recipient mice (Ward et al., 2020) to evaluate the role of the gut microbiota in regulating CNS pathology and disease post-T cell priming. To do this, we generated a pool of encephalitogenic PLP-primed Th17 cells in specific pathogen-free (SPF) female SJL/J donor mice for adoptive transfer into SPF controls or microbiota-depleted SJL/J recipients (**Fig. 1A**). Microbiota depletion was achieved by administration of a broad-spectrum antibiotic cocktail (ABX; vancomycin, neomycin, ampicillin, metronidazole) for 7 days prior to and throughout passive EAE induction. A reduction in total fecal DNA and detectable levels of microbial DNA was confirmed by 16S rRNA sequencing on the same day as EAE induction (**Fig. 1B**). Time to disease onset and incidence were similar between vehicle and ABX-treated recipient mice, indicating that adoptively transferred, pre-primed T cells do not rely on microbiota-derived signals of the host to initiate autoimmune neuroinflammation (**Fig. 1C**). In contrast, later stages of disease were significantly affected. Peak disease scores were lower, and physical mobility was improved at experimental endpoint in microbiota-depleted hosts, leading to an overall reduction in disease burden compared to controls (**Fig. 1D, E**).

**Figure 1.**
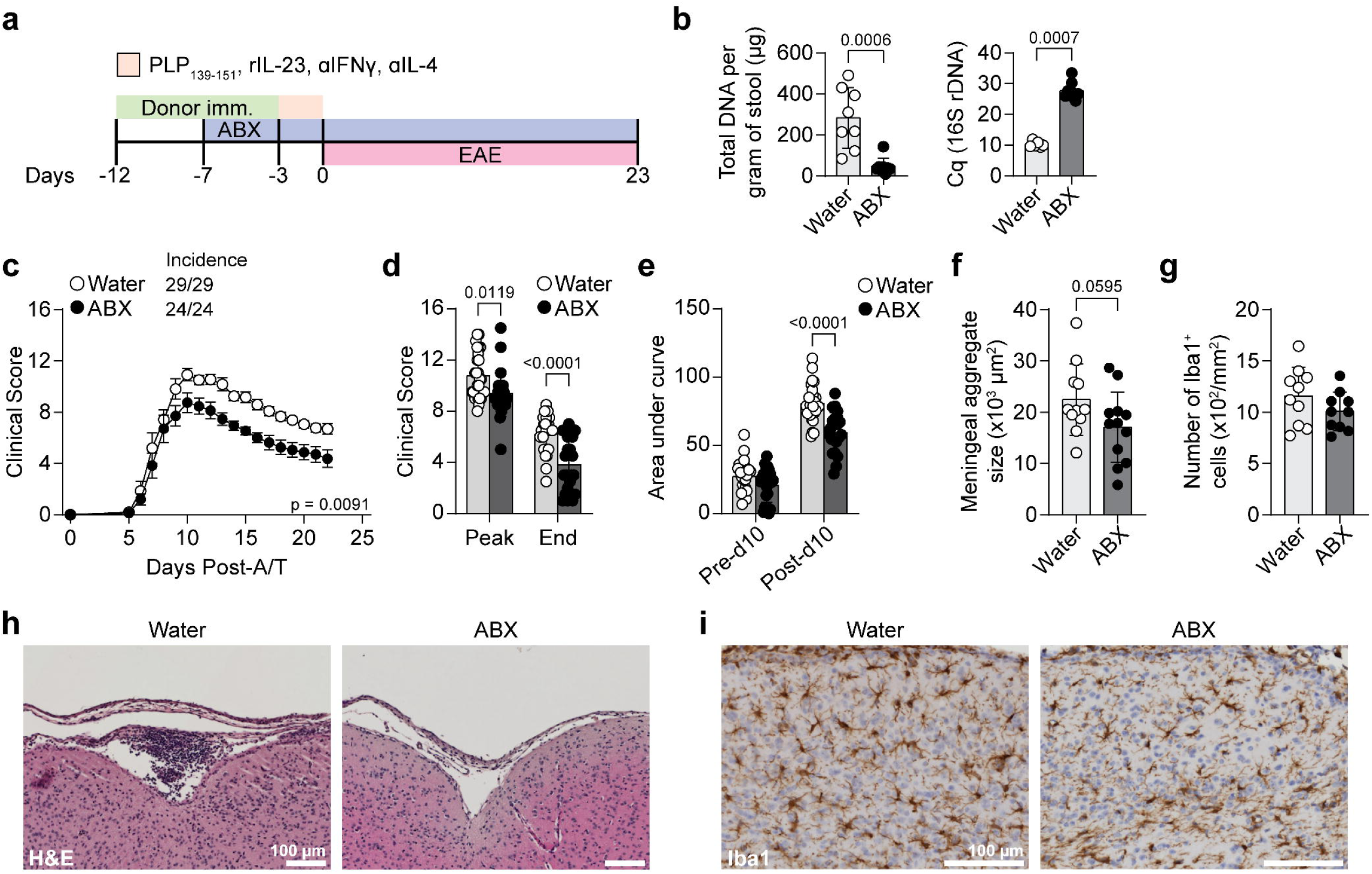
EAE severity is microbiota-dependent. (a) Timeline of donor immunization, ABX treatment, and EAE in young mice (8-12 weeks). (b) Quantification of total DNA in fecal material (left) and Cq values from qPCR detection of 16S rDNA (right). Wilcox rank sum test. (c) EAE clinical scores of mice given water (supplemented with 5% sucrose) or ABX (supplemented with 5% sucrose) *ad libitum* as described in (a). (d) Comparison of peak and endpoint clinical scores. Two-way ANOVA with Sidak multiple comparisons test. (e) Comparison of area under the curve for EAE days 0-10 (inclusive) or from day 11 to endpoint. Mice euthanized before the experimental endpoint were excluded. (f) Quantification of the average meningeal cell aggregate area by H&E staining. (g) Quantification of the average number of Iba1^+^ cells associated with meningeal aggregates. Two-way ANOVA with Sidak multiple comparisons test for (d) and (e). For (f) and (g), Wilcoxon rank sum test. (h) Representative H&E staining images showing meningeal cell aggregates in EAE. (i) Representative Iba1 staining images showing accumulation of Iba1^+^ cells adjacent to meningeal cell aggregation. Scale bars in (h) and (i) represent 100 µm.

Consistent with reduced clinical disease, ABX-treated SJL/J recipients had a modest reduction in the size of meningeal immune aggregates (mean TLT area), but there was no difference in the accumulation of IBA1^+^ myeloid cells adjacent to TLTs compared to mice with an intact microbiota (**Fig. 1F-I**). Collectively, these data indicate that, in a model where encephalitogenic T cells are primed in microbiota-sufficient (no ABX, SPF) donor mice, the microbiome of recipient hosts actively regulates EAE disease outcomes and leptomeningeal neuroinflammation.

### The microbiome of SJL/J mice is minimally affected by chronological age

Aged C57BL/6 and SJL/J mice both develop more severe and non-remitting paralysis following adoptive transfer of encephalitogenic T cells (Atkinson et al., 2022; Zuo et al., 2022). However, the pathology of C57BL/6 mice is primarily in the spinal cord, whereas the cortical grey matter injury and persistent TLT in the brain leptomeninges near areas of microgliosis seen in aged SJL/J mice are hallmarks of progressive MS. Age-associated microbiome shifts have been reported in in C57BL/6 mice (Binyamin et al., 2020; Boehme et al., 2021; D’Amato et al., 2020; Langille et al., 2014), and have been ascribed functional roles in age-associated immune dysfunction (Thevaranjan et al., 2017). However, less is known about the impact of aging on the SJL/J intestinal microbiome. Thus, we investigated three rearing and aging schemes described in the literature with varying levels of environmental control across two independent vivaria (**Fig. 2A**). First, young mice were imported from a commercial vendor and aged in-house. The microbiota of these aged cohorts was compared to young mice recently imported from the same vendor (vendor-sourced, VEN). Second, microbiota samples were compared from age-disparate mice derived from an SJL/J colony bred and raised in-house but not necessarily by the same dam (IH). Third, we compared litters birthed from the same in-house dam (SD) multiple generations apart. In comparison to C57BL/6 mice, SJL/J mice have a shorter life expectancy (Yuan et al., 2009), and we have demonstrated that the age-associated susceptibility to progressive MS-like disease in SJL/J mice begins at approximately 6 months of age and is consistently reproducible at 8 months of age and beyond (Zuo et al., 2022). As such, ‘young’ cohorts were between 8-16 weeks, and ‘aged’ cohorts were >8 months.

**Figure 2.**
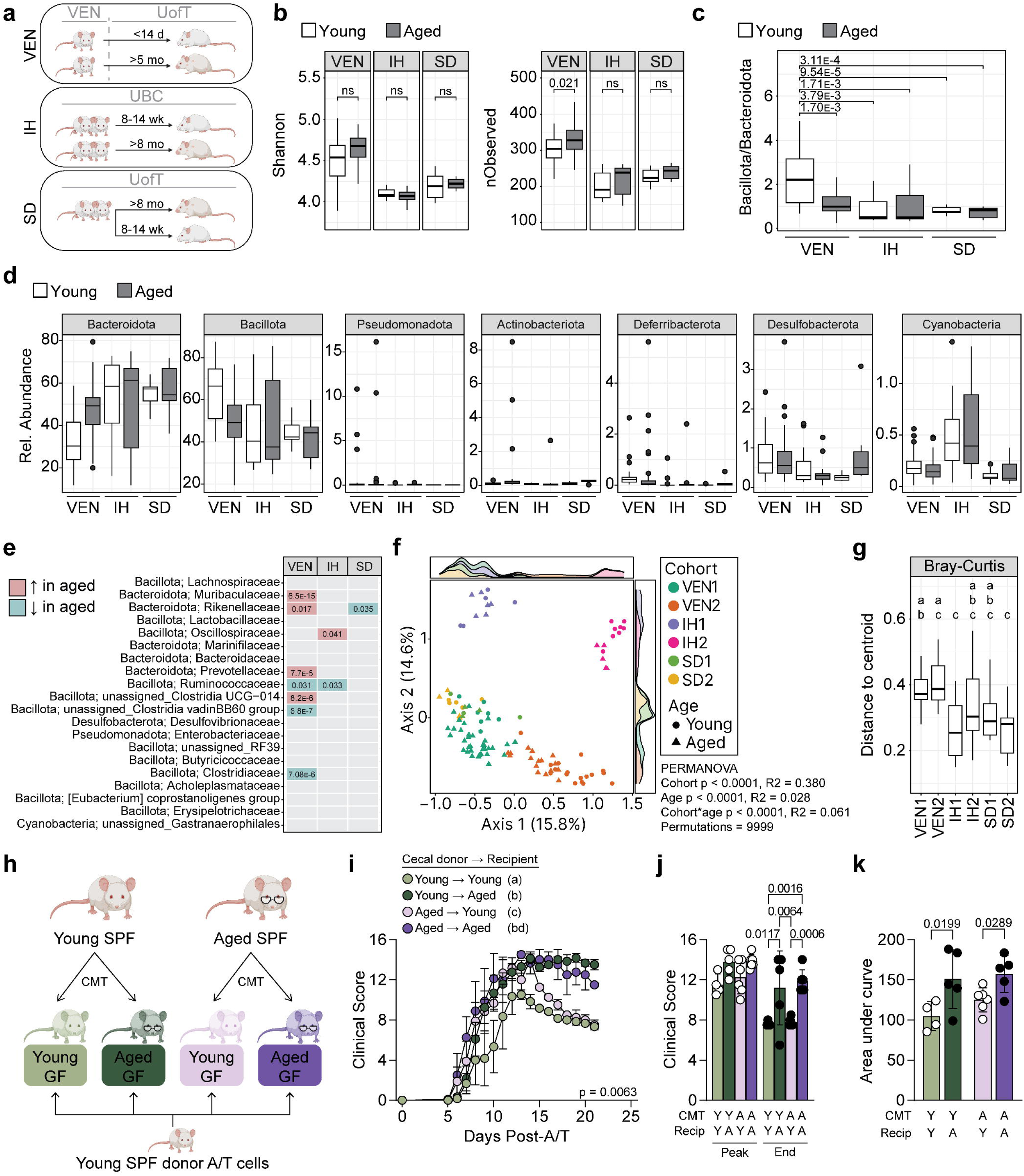
The microbiome of SJL/J mice is minimally affected by biological age. (a) Schematic depicting cohorts of young and aged mice obtained from vendor (VEN), bred within the vivarium but from mixed dams (IH), and bred within the vivarium but from the same breeding pair (SD). (b) Alpha diversity metrics (Shannon diversity, number of observed ASVs) of fecal microbiome from young and aged mice as described in (a). (c) Ratio of Bacillota to Bacteroidota phyla. Kruskal-Wallis test with Dunn multiple comparisons test and Benjamini-Hochberg correction. (d) Comparison of relative abundance of prominent bacterial phyla observed in mice. Kruskal-Wallis test with Dunn multiple comparisons test and Benjamini-Hochberg correction. (e) Comparison of top 20 most abundantly observed bacterial families between young and aged mice within cohorts. Wilcox rank sum test. (f) Principal coordinate analysis (PCoA) plot of Bray-Curtis distances. (g) Comparison of dispersion between combined young and aged mice within cohorts. Kruskal-Wallis test with Dunn multiple comparisons test. Letters indicate shared comparisons (i.e., all groups labeled with “a” are not significantly different). (h) Schematic of FMT experiment. Cecal material were collected and pooled from young or aged SPF donor mice, and transferred to young or aged GF mice. CMT = cecal microbiota transfer (i) EAE clinical scores of young or aged mice given either young or aged mouse donor FMT material. Two-way ANOVA with Tukey multiple comparisons test. Letters indicate shared comparisons. (j) Comparison of peak and endpoint clinical scores. (k) Quantification of area under the curve, excluding mice that were euthanized before the experimental endpoint. For (j) and (k), two-way ANOVA with Sidak multiple comparisons test.

Using 16S rRNA sequencing of fecal pellets to visualize the microbiome community composition of mice in these three husbandry schemes, we observed no significant differences in Shannon diversity or the number of observed amplicon sequence variants (nObs, ASV) between young and aged mice in the most environmentally controlled rearing schemes (IH, SD) (**Fig. 2B**). In contrast, a small increase in the number of observed ASVs was detected in VEN-sourced mice aged in our vivaria compared to recently imported young mice (**Fig. 2B**). Independent of age, the fecal microbiota of VEN mice exhibited significantly higher alpha diversity compared to that of SD and IH groups (**Fig. S1A**).

A diminished Bacillota to Bacteroidota ratio (Blt/Btd) has been reported in studies of human aging and in people with MS (iMSMS Consortium, 2022; Mariat et al., 2009). In the SJL/J cohorts, VEN-derived mice were the only group with a detectable age-associated diminishment of the Blt/Btd ratio. Importantly, the Blt/Btd ratio was highest in young VEN mice compared to all other housing conditions (**Fig. 2C**), suggesting that the reduction in aged cohorts may be due vivarium-based microbiome recalibration rather than age. Although there were inter-cohort differences in abundance of major bacterial phyla (**Table S1**), no age-dependent patterns were detected across all three cohorts (**Fig. 2**). Further, analysis of the 20 most abundant families revealed no consistent differences between young and aged SD, IH, or VEN cohorts that would support a generalized age-dependent effect on the SJL/L microbiome (**Fig. 2E, S1B**).

Beta diversity was assessed by principal coordinate analysis (PCoA) of Bray-Curtis distances (**Fig. 2F**). Differences were detected between young and aged mice in the two independent VEN cohorts, but not within any of the IH or SD cohorts (**Fig. 2F**) and overall, age (or the time spent housed in our vivaria) had a bigger impact on VEN mice compared to the IH and SD cohorts (**Fig. S1C**). Independent of age, the microbiota of SD and IH cohorts had a low distance to the cluster centroid, indicating a high degree of structural similarity among young and aged microbiota (**Fig. 2G, Table S1**). Conversely, the VEN cohorts had a larger dispersion and therefore less uniform microbiome composition than either the IH or SD cohorts.

Collectively, these data indicate that under environmentally controlled conditions, age alone (>8 months) has a minimal effect on the fecal microbiome of laboratory SJL/J mice. Moreover, the comparison of different breeding and housing strategies on aging microbiome composition highlights the importance of experimental design when analyzing microbiota contributions to disease phenotypes.

### The aged SJL/J microbiota is not sufficient to promote non-remitting EAE

Although 16S rRNA-based analysis indicated strong similarities in the microbiomes of young and aged SJL/J mice, functional differences beyond the limit of 16S detection (e.g. low abundance microbes or strain specific differences that alter metabolite production) could contribute to age-dependent EAE outcomes. To assess this possibility, we took advantage of the fact that the host microbiome modulates the chronic phase of T cell transfer EAE. Accordingly, microbiota collected from the cecal contents of young or aged SPF IH-reared donors was transferred (cecal microbiota transplant, CMT) into sex-matched young or aged GF recipients (**Fig. 2H**). No significant differences in alpha diversity were observed in the fecal microbiota across recipient groups (**Fig. S1D**). Notably, the microbiota composition of young CMT → young ex-GF hosts and aged CMT → aged groups shared the least overlap in PCoA space, and Bray-Curtis dissimilarity analysis suggested that age of the recipient (*p* = 0.004) and age of the microbiota donor (*p* = 0.001) can both influence microbial community structure in ex-GF mice (**Fig. S1E, Table S1**). Differential abundance testing revealed a small number of ASVs that were enriched in either young or aged FMT recipients, including several bacterial families with members enriched in both FMT groups (**Table S1**).

We tested the impact of heterochronic murine CMT on T cell adoptive transfer EAE using Th17 cells generated in young SPF mice. Despite the modest shift in microbiome composition following FMT, age-disparate FMTs did not alter EAE outcomes: aged recipients developed chronic disease regardless of the age of the transferred microbiota, and young recipients recovered over time independent of the age of the transferred microbiota (**Fig. 2I**). While peak EAE severity did not differ among groups, the extent of recovery mirrors the expected trajectory of the age of the recipient (**Fig. 2J**). Similarly, the AUC revealed a significant effect of the age of the recipient, but not the FMT age, in terms of overall disease severity (**Fig. 2K**). Collectively, these data indicate that an aged SJL/J microbiome is insufficient to drive chronic disease in young recipient mice.

### Xenografted microbiota from healthy, age-disparate family members can modify EAE outcomes

While we have shown that vivarium-housed SJL/J mice do not exhibit age-dependent microbiome changes, prophylactic antibiotic treatment of mice that subsequently received encephalitogenic Th17 cells indicated that the post-priming phase of disease is microbiota-sensitive in our EAE model. Unlike vivarium-reared mice, age-associated shifts in human microbiomes are influenced by environmental and lifestyle factors (Ghosh et al., 2022b). Thus, we hypothesized that some aged human microbiomes may be sufficient to alter the remission trajectory of young SJL/J mice. If so, this would provide an opportunity to mine for translationally relevant microbiome differences that influence disease progression.

To test this hypothesis, we generated humanized-microbiota mice via FMT from healthy human donors, selecting sex-matched, first-degree relative adult donor pairs living in the same household, who were separated in age by at least 25 years (**Table 2**). The household design has previously been validated to improve statistical power in population microbiome studies and is the current gold-standard to control for environmentally-driven microbiome variation (iMSMS Consortium, 2020; 2022). FMT from four donor pairs were evaluated in young mice that had undergone a microbiota-depleting regimen previously shown to support human FMT (hFMT) engraftment on par with GF recipients (Staley et al., 2017). After 4 weeks of host and microbiota stabilization, we induced EAE via adoptive transfer of PLP-primed encephalitogenic Th17 cells generated in young SPF donor mice (**Fig. 3A**) to examine whether an aged hFMT could promote an “aged” EAE phenotype (i.e., chronic, non-remitting EAE) in chronologically young recipient mice.

**Figure 3.**
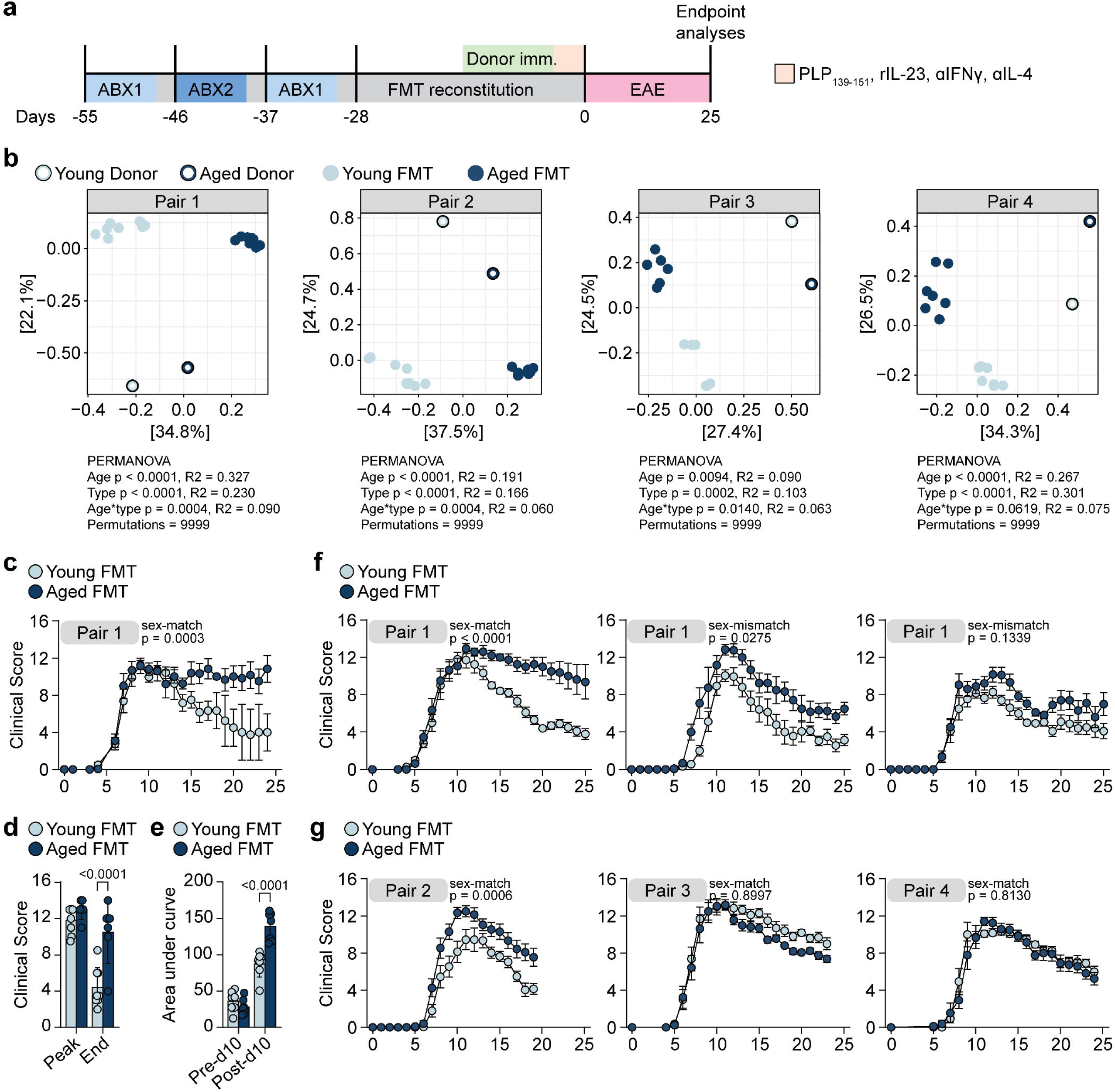
Xenografted microbiota from healthy, age-disparate family members can modify EAE outcomes. (a) Schematic of antibiotic pre-treatment and FMT timeline. ABX1 = vancomycin, neomycin, ertapenem; ABX2 = ampicillin, cefoperazone, clindamycin. (b) PCoA plot of Bray-Curtis distances among mice given FMT from donor pair 1. (c) EAE clinical scores of young mice given FMT from donor pair 1. (d) Comparison of peak and endpoint clinical scores. Two-way ANOVA with Sidak multiple comparisons test. (e)(g)(i) EAE clinical scores of young mice given FMT from donor pairs 2 through 4. Data for each donor pair are from one representative experiment of 2 total (1 at UofT, 1 at UBC). (f)(h)(j) PCoA plots of Bray-Curtis distances among mice given FMT from donor pairs 2 through 4. 16S data from donor pair 1 derived from fecal samples collected from UofT experiments, and donor pairs 2-4 from both UBC and UofT experiments. (k) Experimental repeats conducted at UBC vivarium (left, middle) and UofT (right). For all EAE clinical score curves, two-way ANOVA with Tukey multiple comparisons test.

Each donor pair was independently evaluated at both UofT and UBC sites to minimize confounding vivarium-specific effects. The fecal microbiome of hFMT recipients revealed that each pair of FMT donor materials elicited distinct bacterial communities in recipient mice (**Fig. 3B**). The hFMT from the aged donor of Pair 1 exacerbated EAE and blunted remission compared to the young hFMT, a result that was consistent across multiple repeats at both vivaria (**Fig. 3C-F**). Notably, the effect of the aged hFMT was more robust when the human FMT donors and recipient mice were sex-matched (**Fig. 3C, F**). Thus, all subsequent young and aged hFMT were used to colonize mice sex-matched to the donor pairs. The aged hFMT from Pair 2 also exacerbated EAE severity chronicity compared to the young counterpart, while there were no differences in EAE outcomes between the young and aged hFMT from the other two pairs (**Fig. 3G**). As we did not pre-screen donors based on microbiota-modifying lifestyle factors (e.g., diet and exercise), substantial variability between donor pairs is not unexpected. Collectively, these data demonstrate that hFMT from some otherwise healthy aged donors can exacerbate passive EAE in SJL/J mice, providing us with communities for testing age-associated metabolites that impact the post-priming phase of disease.

### Heterochronic microbiota xenografts confer structurally distinct communities and influence EAE-induced neuroinflammation

To investigate potential drivers of disease enhancement, we further examined the microbiome composition and pathophysiology of mice colonized with hFMT from donor Pair 1, which had the most striking difference in clinical disease course. Differential abundance testing of fecal microbiomes of Pair 1 hFMT colonized mice revealed a total of 53 significantly altered taxa (*p* < 0.00001) (**Fig. 4A, Table S1**). Taxa significantly enriched in recipients of the aged hFMT include *Fusobacterium ulcerans*, *Parabacteroides merdae*, and two distinct *Ruminococcus* species belonging to the *R. gnavus* or *R. torques* groups. Conversely, taxa significantly higher in young hFMT mice include *Akkermansia muciniphila*, an unspecified *Roseburia* species, *Sutterella wadsworthensis* as well as another unspecified *Sutterella* species, and *Adlercreutzia equolifaciens* (**Fig. 4B**). Therefore, although all recipient mice tested were young, introduction of an aged human microbiome via FMT can elicit severe disease accompanied by alterations in fecal microbiome community structure.

**Figure 4.**
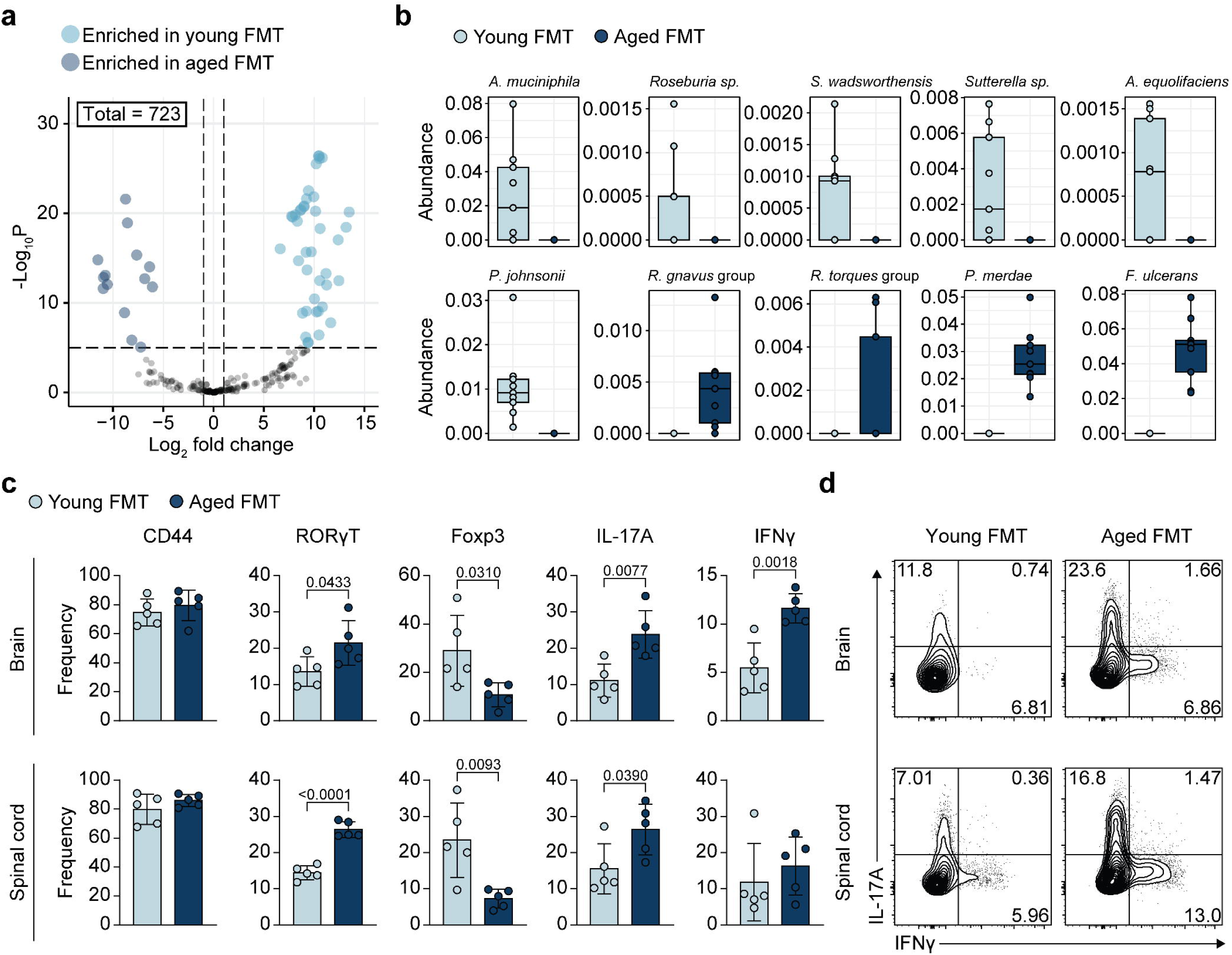
Heterochronic microbiota xenografted microbiota confer structurally distinct communities and influences EAE neuroinflammation. (a) Volcano plot summarizing differential abundance analysis conducted using the DESeq2 package (Love et al., 2014). Log_2_FC value cutoff shown in volcano plot = 1. (b) Direct comparisons of selected taxa differentially abundant in either young or aged FMT recipient mice. (c) Flow cytometric analysis of brain and spinal cord-infiltrating CD4^+^ T cells during acute peak phase in mice given FMT from donor pair 1. Wilcoxon rank sum test. (d) Representative flow cytometry plots of brain and spinal cord-derived CD4^+^CD44^+^ T cells.

Next, we assessed the phenotype and cytokine-producing potential of CNS-penetrating T cells at the peak phase of EAE. We hypothesized that differences in the inflammatory milieu of the CNS at this timepoint would reflect the subsequent disease trajectory (i.e. remission or chronic paralysis). The frequency of pathogenic Th17 cells RORγT^+^ was elevated in the brain and spinal cord of aged hFMT colonized animals, which was associated with a corresponding decrease in CD4^+^CD44^+^ Foxp3^+^ T regulatory (Treg) cells (**Fig. 4C**). Consistent with these observations, the proportion of PLP_139-151_-responsive IL-17A- and IFNγ-producing CD4^+^ T cells were markedly increased in the CNS of mice colonized with the aged hFMT (**Fig. 4D**). Collectively, these data demonstrate that the aged hFMT microbiota-dependent shift toward non-remitting EAE is associated with an accumulation of neuroinflammatory T cell populations known to promote SJL EAE and MS pathogenesis.

### Microbiota-dependent Indole-3-propionic acid (IPA) promotes EAE resolution

We postulated that the exacerbated EAE severity and chronicity in aged hFMT colonized mice could be mediated via microbiota-derived metabolites that modulate host cells involved in sustaining neuroinflammation or neurodegeneration. To test this, we collected serum and the cerebral cortex of mice harboring the remission-prone ‘young’ or non-remitting ‘aged’ hFMT from donor Pair 1 at four weeks post-FMT for untargeted metabolomic analysis. We identified 53 over- and 31 under-represented metabolites in the cortex of young hFMT colonized mice compared to aged hFMT cortex, and 39 and 37 metabolites that were over- or under-represented in the serum of mice harboring the young hFMT compared to aged hFMT recipients (**Fig. 5A, Table S2**). Hits of interest in the serum included components of soy isoflavone metabolism, dihydrodaidzein, equol, and glycitein 4’-O glucuronide, and indole-3-propionic acid (IPA), a derivative of tryptophan produced by gut microbiota (**Fig. 5B**). Other serum-derived metabolites included an Arg-Leu-Val tripeptide and 2-ethylpyridine (**Fig. 5B**). In the cortex, mice colonized with the aged hFMT had lower levels of n-acetyl-tyrosine and increased levels of a Gly-Leu dipeptide (**Fig. 5C**), together suggesting potential effects on dopamine metabolism or neurotransmission.

**Figure 5.**
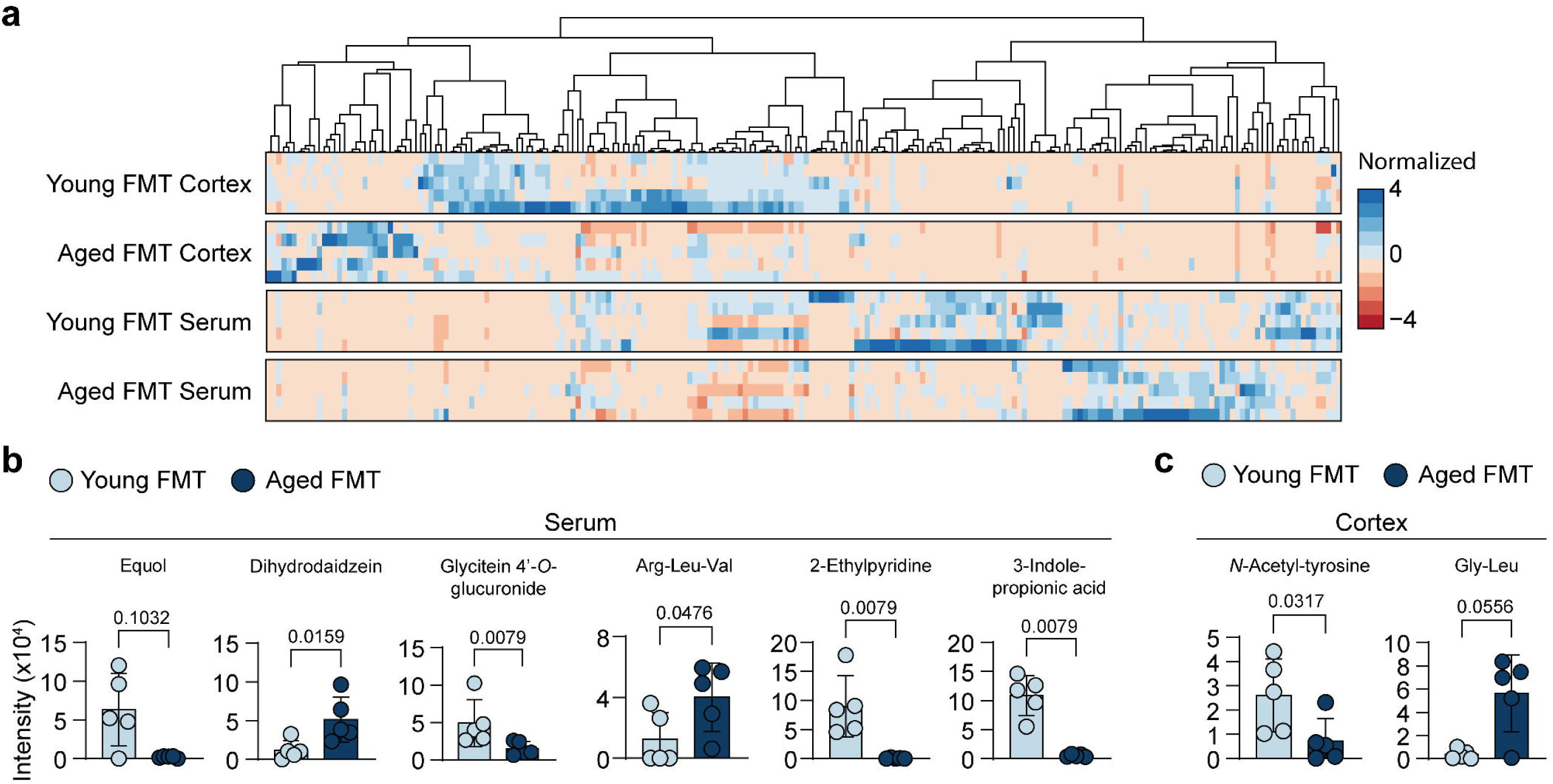
Differential metabolomic profiles of mice with microbiome xenografts. (a) Heatmap of all detected compounds in brain cortical tissue and serum of mice given FMT from donor pair 1. Values are normalized internally between samples. (b),(c) Highlighted differentially abundant metabolites enriched in either serum (b) or cortical tissue (c) of FMT recipient mice. Wilcoxon rank sum test.

We focused our analysis on metabolites derived from bacterial metabolism that were consistently present or absent in mice colonized with young vs aged hFMT. One specific metabolite fulfilling these criteria was IPA, as it was nearly undetectable in the serum of aged hFMT recipients but highly abundant in recipients of the young hFMT (**Fig. 5B**). Since tryptophan and several of its derivative metabolites, including indole compounds, have been previously implicated to have potent immunomodulatory and EAE-ameliorative roles (Fettig and Osborne, 2021; Rothhammer et al., 2016; Rouse et al., 2013), we hypothesized that the lack of IPA in the serum of aged hFMT recipient mice could contribute to exacerbated EAE. To test this, chronologically young mice were microbiota-depleted and colonized with the aged hFMT from Pair 1. After 4 weeks of microbiota reconstitution, mice were treated with daily injections of IPA (or vehicle control) for 7 days prior to Th17 cell adoptive transfer EAE and throughout the course of clinical disease (**Fig. 6A**). Consistent with the untargeted metabolomic screening results, IPA was undetectable in the serum of vehicle-treated mice, but circulating IPA levels were elevated in IPA-supplemented mice at the time of EAE induction (**Fig. 6B**). IPA treatment had no effect on EAE susceptibility (incidence was similar between IPA and control groups) (**Fig. 6C**). However, while vehicle treated mice developed severe, non-remitting EAE, IPA supplementation promoted rapid clinical remission (**Fig. 6C-E**).

**Figure 6.**
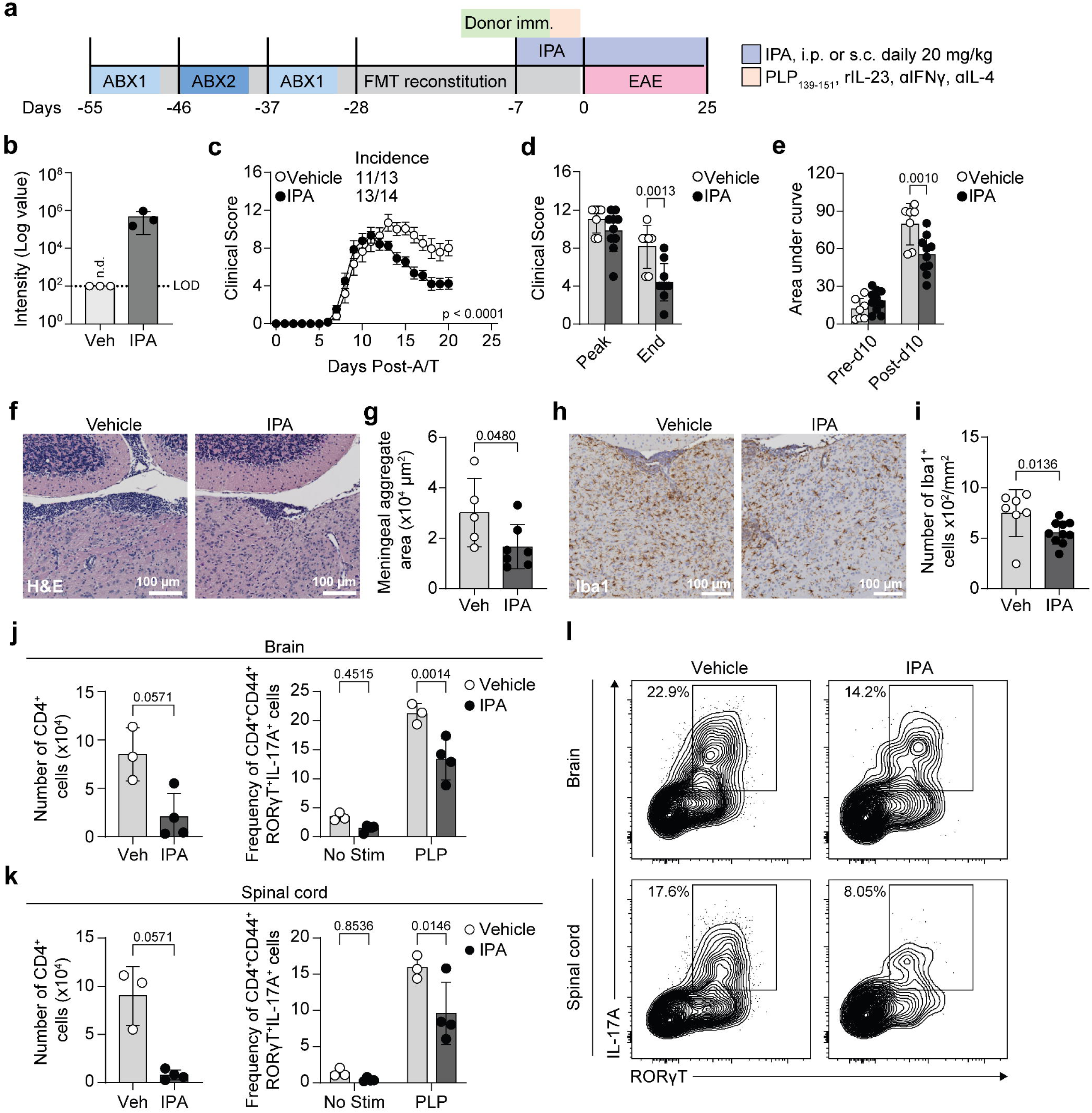
IPA drives resolution of clinical EAE and inhibits T cell IL-17A production. (a) Schematic of experimental setup for antibiotic pre-treatment, FMT reconstitution, and IPA treatment. (b) Quantification of serum IPA levels prior to A/T. LOD = limit of detection. n.d. = not detected. (c) EAE clinical score of FMT recipient mice treated with IPA or vehicle. (d) Peak and endpoint clinical scores, excluding mice that were euthanized prior to the experimental endpoint. (e) Area under the curve of EAE day 0-10 (inclusive) or from day 11 to the experimental endpoint. (f) Representative images for H&E staining. (g) Quantification of meningeal aggregate area. (h) Quantification of meningeal aggregate-adjacent Iba1^+^ cells. (i) Representative images for Iba1^+^ cells. (j),(k) Flow cytometric analysis, total number of CD4^+^ cells and frequency of CD4^+^CD44^+^ IL-17A-producing cells obtained from the brain (j) and spinal cord (k). (l) Representative flow cytometry plots for (j) and (k). Only PLP-stimulated samples are shown. Two-way ANOVA with Tukey multiple comparisons test for (c). Wilcoxon rank sum test for (g), (i), left panels of (j) and (k). Two-way ANOVA with Sidak multiple comparisons test for (d), (e), and right panels of (h) and (i).

Reflecting the reduced clinical severity in IPA-supplemented aged hFMT animals, average TLT size and TLT-associated Iba1^+^ cell counts were both reduced in the cerebral cortex of IPA-treated mice (**Fig. 6F-I**). Further, absolute numbers of CD4^+^ T cells in the CNS of IPA-treated mice were reduced compared to vehicle controls (**Fig. 6J left, K left**). Indole-mediated immunosuppression has previously been described to impair Th17 activation (Rouse et al., 2013). Consistent with this, activated CD4^+^CD44^+^ T cells isolated from the CNS of IPA-treated mice exhibited reduced IL-17A production following *ex vivo* restimulation with PLP_139-151_ (**Fig. 7J-L**). Overall, these data suggest that microbiota-dependent IPA production restrains neurodegenerative potential in a mouse model of progressive MS, and that in an IPA-deficient setting, exogenous IPA supplementation can suppress Th17-associated neuroinflammation.

**Figure 7.**
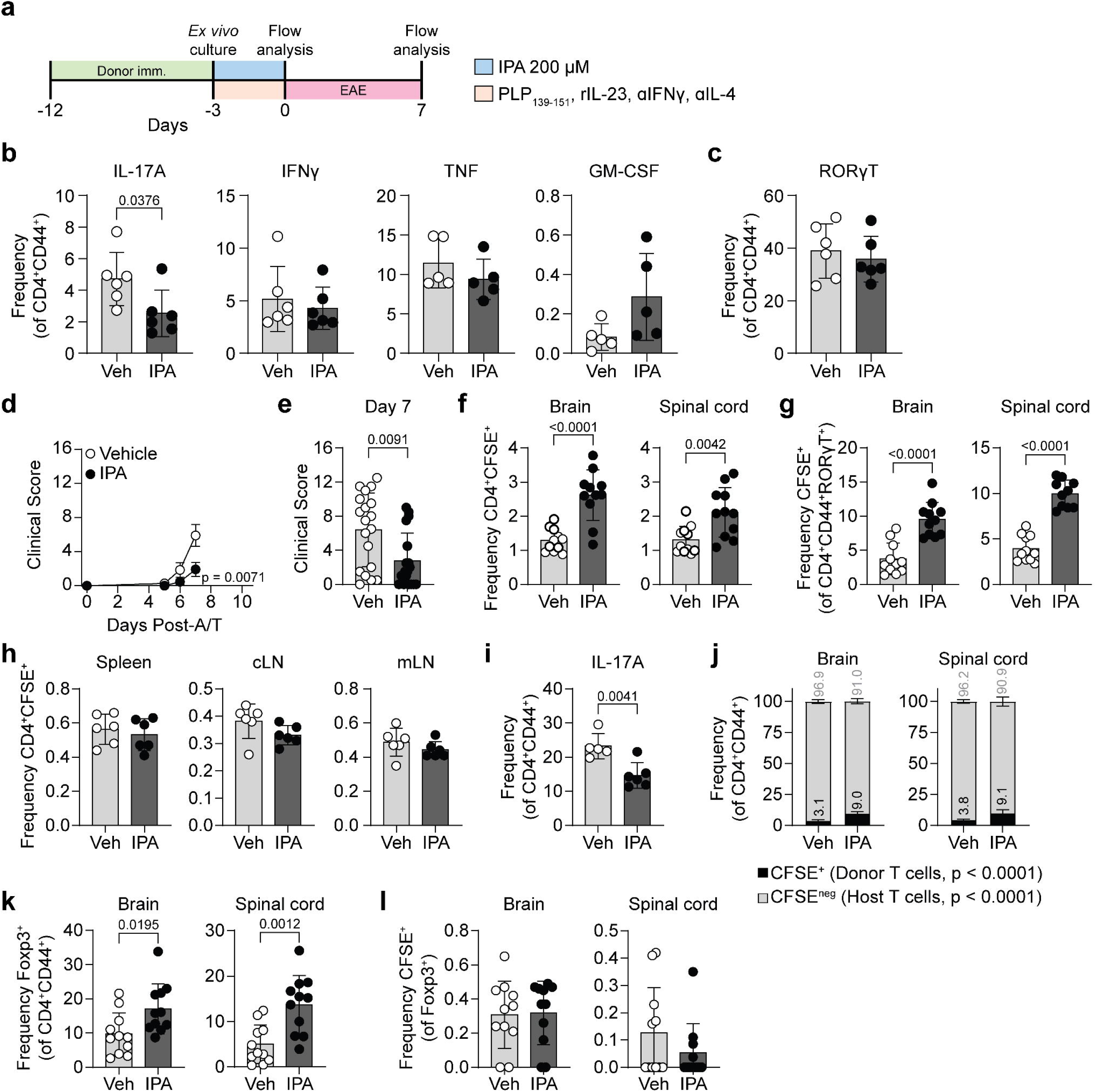
IPA reduces IL-17A producing capacity and encephalitogenic potential of A/T cells. (a) Schematic of experimental setup for IPA treatment of Th17-polarized cells *in vitro*. (b) Flow cytometric analysis of IL-17A, IFNγ, TNF, and GM-CSF in Th17 cells treated with IPA or vehicle. (c) Frequency of RORγT-expressing CD4^+^ T cells with IPA treatment. (d) EAE clinical scores of mice given donor cells pre-treated with IPA or vehicle. (e) Clinical score at day 7 post-A/T in mice given donor cells pre-treated with IPA or vehicle. (f) Flow cytometric analysis of frequency of total CFSE^+^ CD4^+^ T cells in the brain (left) and spinal cord (right) at day 7 post-A/T. (g) Frequency of CFSE^+^ CD4^+^CD44^+^RORγT^+^ cells in the brain (left) and spinal cord (right) at day 7 post-A/T. (h) Frequency of CFSE^+^ CD4^+^ T cells in the spleen, cervical lymph nodes (cLN), and mesenteric lymph nodes (mLN) of mice 7 days post-A/T. (i) IL-17A^+^ frequency of CD4^+^CD44^+^ T cells in the brain 7 days post-A/T. (j) Bar plots showing frequencies of CFSE^+^ (donor-derived) and CFSE^neg^ (recipient-derived) CD4^+^CD44^+^ T cells in the brain (left) and spinal cord (right) of mice 7 days post-A/T. Significance shown applies to both brain and spinal cord. (k) Frequency of total Foxp3^+^ CD4^+^ T cells in the brain (left) and spinal cord (right) 7 days post-A/T. (l) Frequency of Foxp3^+^CFSE^+^ cells in the brain (left) and spinal cord (right) 7 days post-A/T. Wilcoxon rank sum test for (b), (c), (e), (f), (g), (h), (i), (k), (l). Two-way ANOVA with Tukey multiple comparisons test for (d), (j).

### Indole-3-propionic acid (IPA) reduces T cell IL-17A production and encephalitogenic potential

In the adoptive transfer SJL/J model, EAE initiation is dependent on the effector activity of transferred encephalitogenic Th17 cells, but subsequent waves of endogenous T cell activation and CNS recruitment are critical for disease chronicity (Pikor et al., 2015). When EAE was induced by encephalitogenic Th17 cells derived from SPF mice, *in vivo* IPA supplementation did not affect EAE susceptibility but led to subsequent EAE remission and reduced accumulation of Th17 cells in the CNS, suggesting that IPA may restrain the secondary waves of autoreactive T cells that promote non-remitting EAE.

To directly test the effects of IPA on encephalitogenic Th17 cell priming, we isolated leukocytes from PLP-immunized donor animals and cultured the cells *ex vivo* under Th17-skewing conditions in the presence of PLP_139-151_ and IPA or vehicle (0.1% DMSO) (**Fig. 7A**). After 72 hours, the frequency of CD4^+^CD44^+^ T cells expressing IL-17A (but not IFNγ, TNF, or GM-CSF) was reduced by the presence of IPA (**Fig. 7B**). No changes in the expression of RORγT were observed (**Fig. 7C**), indicating a reduction in Th17 effector function rather than polarization. IPA exposure had no detectable effect on Treg or Th1 differentiation or general T cell activation, as indicated by expression of Foxp3, Tbet, CD69, and CXCR6, respectively (**Fig. S2A**).

Next, we assessed the effects of IPA treatment during *ex vivo* encephalitogenic Th17 cell priming on EAE susceptibility and the endogenous T cell compartment of young, conventional recipient mice. Due to the lack of a congenic line of SJL/J mice, adoptively transferred donor T cells were fluorescently labelled with CFSE to facilitate tracking in the recipients, as we have previously described (Florescu et al., 2025). Consistent with the requirement for IL-17A to initiate the pathogenic cascade (Pikor et al., 2015), mice injected with IPA-treated cells exhibited slightly delayed disease onset, with fewer mice exhibiting paralytic symptoms at day 7 post-A/T (clinical score > 2; 13 of 19 mice in vehicle group, 6 of 18 mice in IPA-treated group) (**Fig. 7D, E**). In order to quantify and phenotype CFSE^+^ donor T cells, tissues were harvested at day 7. Despite the reduced signs of clinical disease, the frequency of donor-derived CFSE^+^CD4^+^ T cells and CFSE^+^RORγT^+^ Th17 cells was significantly higher in the CNS of mice that received IPA-treated donor cells (**Fig. 7F, G**). The relative enrichment of CFSE^+^ donor T cells was only observed in the CNS, not in peripheral tissues (**Fig. 7H**), indicating that IPA does not impair initial Th17 CNS recruitment. However, the frequency of brain-infiltrating CD4^+^CD44^+^ T cells (combined CFSE^+^ and CFSE^neg^) expressing IL-17A was reduced in IPA-A/T recipients (**Fig. 7I**), suggesting that IPA may dampen the ability of autoreactive T cells to recruit neuroinflammatory endogenous T cells. We used CFSE^neg^ as a proxy marker of host-derived cells to address this question. This strategy suggested that the fraction of host-derived (CFSE^neg^) cells in the pools of activated CD44^+^ CD4^+^ T cell and RORγt^+^ Th17 cell populations was more substantial when EAE was induced with vehicle-treated Th17 cells than with IPA-treated Th17 cells (**Fig. 7J**). Notably, IPA exposure during T cell priming led to a relative enrichment of Foxp3^+^ Treg in the CNS at day 7 post-EAE induction (**Fig. 7K**). The frequency of CFSE^+^ donor cells was negligible in both vehicle and IPA-A/T recipients (**Fig. 7L**), which is consistent with the donor T cell population coming from a Th17 skewed culture, and suggests that CNS Tregs are host-derived. Together, these data highlight the impaired ability of the IPA-treated encephalitogenic T cells to promote second wave recruitment of endogenous pathogenic Th17 cells into the CNS, providing a potential mechanism of action for microbiota-derived IPA in limiting progressive MS-like disease.

## Discussion

In this study, we demonstrate that the role of the microbiota in EAE susceptibility and disease trajectory can be independently assessed when encephalitogenic T cells are primed in a microbiota-sufficient environment and transferred into host mice with an altered microbiome. Heterochronic mouse-to-mouse CMT demonstrated that an aged mouse microbiome was insufficient to confer the progressive MS-like phenotype associated with chronological age in young SJL/J mice and vice versa. In contrast, FMT of microbiota from some aged human donors was able to confer an aged EAE phenotype in young mice. Chronologically young recipient mice colonized with an aged human donor microbiota that reliably induced an aged EAE phenotype demonstrated significant alterations in the metabolome of their blood and cortex, and this pipeline identified IPA as a modulator of neuroinflammation. Collectively, these results provide new insights into limitations of using mouse FMT for aging studies and demonstrate that applying human FMT to our novel EAE model provides a mechanism for unveiling metabolites that alter disease progression.

Many reports investigating the effect of the gut microbiome on neuroinflammation use models that involve immunization against myelin peptides to induce EAE (Miyauchi et al., 2020; Ochoa-Repáraz et al., 2009; Seifert et al., 2018). However, due to effects of the microbiota on systemic immune priming, this approach confounds specific examination of how the microbiota modulates primed cell effector activity in the CNS. The A/T EAE approach allows the distinction of the priming and effector phases of EAE. Focusing on a model that elicits grey matter pathology relevant to MS progression (Ward et al., 2020; Zuo et al., 2022), we find that microbiota-depletion in recipient, but not donor mice, impacts peak and post-peak disease severity but not onset or incidence of EAE. Our results are consistent with those from an adoptive T cell transfer EAE model in rats that indicate the lung microbiome regulates EAE outcomes by modulating CNS-intrinsic microglia activity (Hosang et al., 2022). Collectively, these data suggest that there is a significant role for microbial regulation of immune cell infiltration and activity in the CNS independent of the initial priming event.

Multiple factors must be considered when conducting microbiome studies in mice (Goodrich et al., 2014; Macpherson and McCoy, 2015). The major determinants of murine intestinal microbiota composition are vertical inheritance, vivarium-dependent factors such as diet, water acidification, housing and handling, and genetic differences among strains or genetically-modified strains (Ericsson et al., 2021, 2018, 2015; Guo et al., 2022; Long et al., 2021; Moeller et al., 2018; Parker et al., 2018; Rasmussen et al., 2019; Stappenbeck and Virgin, 2016). Furthermore, vendor-to-vendor differences in microbiome composition and strain genetics are notorious for their impact on the immune response and by extension, on disease models such as EAE (Ivanov et al., 2009; Summers deLuca et al., 2010). When these factors were controlled to the best of our ability, we did not observe a major impact of age on microbiome composition or diversity in SJL/J mice over 8 months of age. While we observe an age-dependent shift in the EAE clinical phenotype in SJL/J mice at this timepoint and thus selected 8-12 month-old mice for our analyses, we also cannot discount the possibility that assessing mice at more extreme ages could reveal taxa-level or functional differences in the microbiota. For SJL/J mice, this presents further challenges due to the shortened longevity and higher incidence of spontaneous lymphomas (Chow and Ho, 1988; Yuan et al., 2009).

Alternatively, humans experience a wide variety of environmental exposures that modify gut microbiota which are absent in the vivarium (e.g. dietary changes, infection history, circadian rhythm disruptions, anthropogenic pollutants, antibiotic exposure, and other lifestyle choices). Additionally, the conventional laboratory mouse gut microbiome is notoriously simple, is taxonomically dissimilar to the human gut microbiome, and is relatively less resilient (Ng et al., 2019). Thus, FMT from human donors has the capacity to introduce some of the complexities and diversity that arise in the microbiome over the course of a lifetime. Our approach of recruiting pairs of same-sex, first-degree relatives that share a household limits the genetic and environmental confounders between them and allows greater confidence in identifying microbes and their associated metabolites that may be altered by age (iMSMS Consortium, 2020). Using this design, we identified differentially abundant taxa present in hFMT donor material that are associated with exacerbated disease. Whether these changes are due to age alone is still under speculation, as we did not collect dietary or lifestyle information and only pre-screened donors based on general daily health, nor is our study sufficiently powered or designed to identify significant differences with age among our human donors.

Some key taxa differences that may contribute to FMT-driven EAE phenotypes include the high abundance of *Akkermansia muciniphila* detected in recipients of the remission-prone young hFMT from donor Pair 1. *A. muciniphila* has been suggested to promote tolerogenic immune functions in multiple EAE studies (Cox et al., 2021; Gandy et al., 2019), suggesting that its presence may be one mechanism promoting EAE remission. However, others have suggested that this protection can be strain-dependent (Schwerdtfeger et al., 2025), which may explain why *A. muciniphila* has, in other cases, been shown to promote a proinflammatory T cell phenotype (Cekanaviciute et al., 2017). We found two *Sutterella* species, including *S. wadsworthensis*, to be decreased in aged hFMT compared to young – a similar pattern was found to be reduced in an MS cohort (Miyake et al., 2015). Short-chain fatty acid producers such as *Roseburia* species were also enriched in young hFMT mice. Consistent with datasets from elderly individuals (Wang et al., 2019; Yan et al., 2022), the relative abundance of *Parabacteroides merdae* and *Ruminococcus gnavus* was elevated in the fecal microbiome of mice colonized with the aged FMT. *Fusobacterium ulcerans* was the top enriched taxa identified in the aged hFMT, however information on its relevance to either aging or MS are currently lacking. Interestingly, the related species *F. nucleatum*, a periodontal pathogen, has been linked to higher disability scores in people with MS (Naito et al., 2025), but further work is needed to determine whether *F. ulcerans* has similar disease-promoting properties.

Further, we found that the different microbial communities introduced into experimental mice conferred distinct metabolomic profiles in the serum and cortex. Several metabolites have been identified as particularly enriched in either the young or aged FMT mice, some of which have specific relevance to the gut-brain axis. These were indole-3-propionic acid (IPA), equol, and dihydrodaidzein. IPA is one of several indole derivatives that is produced solely by tryptophan metabolism by bacteria. This family of indole compounds have already been implicated in both CNS and non-CNS diseases, but IPA in particular has been shown to be important in maintaining gut barrier integrity and exhibit immunomodulatory activity, possibly through either the pregnane X receptor or aryl hydrocarbon receptor (AhR) (Huang et al., 2022; Rothhammer et al., 2016; Venkatesh et al., 2014). Furthermore, age-associated reductions of serum tryptophan and indole levels have been reported in both mice and humans (Ruiz-Ruiz et al., 2020; Wu et al., 2021). We also report here that a likely mechanism by which IPA is protective in our EAE model is the reduction of IL-17A production by inciting encephalitogenic Th17 cells. This is consistent with studies using other indole derivatives, such as indole-3-lactic acid (Wilck et al., 2017) and indole-3-carbinol (Rouse et al., 2013). Given the evidence that these pathogenic T cells can become “licensed” in the gut environment prior to infiltrating the CNS and the abundant evidence linking gut microbial regulation of Th17 cells, the gut likely serves as a secondary site for receiving additional polarization signal and may alter the encephalitogenic potential of Th17 cells before their migration into the CNS. Interestingly, independent studies have also shown that certain indole derivatives can promote, rather than inhibit Th17 function (Rebeaud et al., 2025). Although these compounds share activity through the AhR receptor, there is evidence that different AhR agonists can have divergent effects (Quintana et al., 2008) in polarizing T cells. The range of effects elicited by various indole compounds on T cell populations, especially Tregs and Th17, has yet to be fully elucidated.

Equol and dihydrodaidzein are both derived from intestinal processing of isoflavones found in legumes exclusively by gut bacteria, and their reciprocal abundance in young and aged FMT mice may indicate a deficiency in the ability of the gut microbiota to process dihydrodaidzein into equol. The functional implications of the lack of equol in EAE are relatively understudied, however equol has been reported to be neuroprotective *in vitro* (Subedi et al., 2017) and an isoflavone-rich diet ameliorates EAE severity (Jensen et al., 2021). This protection depended on the presence of known isoflavone-metabolizing bacterium *Adlercreutzia equolifaciens*. Of note, EAE in *A. equolifaciens*-monocolonized mice was exacerbated when the diet lacked isoflavone, suggesting an important role of diet (Jensen et al., 2021). Interestingly, our 16S sequencing efforts also identified an enrichment in *A. equolifaciens* in yFMT mice (Maruo et al., 2008). *Adlercreutzia* has been reported to be reduced in people with MS, suggesting a loss of neuroprotective or anti-oxidative stress mechanisms (Chen et al., 2016b). The identification of several putative EAE regulators from a single FMT donor pair highlight the utility of this model for identifying therapeutically-relevant microbiome-derived molecules. Similar to characterization of the remission-promoting role of IPA presented here, follow-up studies could test the role of equol supplementation on EAE initiation, severity, and clinical trajectory.

There are inherent technical factors that may impact the consistency of microbiome-related findings across studies. The processing of the donor material, the number of “doses” the recipient mice are exposed to, and the recipient gut environment (method of clearing the endogenous bacterial niche), are known to impact donor material quality and influence microbiota engraftment (Costea et al., 2017; Gopalakrishnan et al., 2021; Le Roy et al., 2019). In our study, we aimed to standardize methods across vivariums whenever feasible, and the EAE phenotypes observed in multiple hFMT donors were highly reproducible across independent investigators at geographically separated vivariums. Nonetheless, we cannot exclude the possibility that other FMT protocols may impact the EAE phenotypes differently.

There are some limitations to our study. Most studies in C57BL/6 mice consider 16 months or higher to be aged with pronounced changes in a number of physiological and biomarker metrics (Xie and Ehninger, 2023). However, due to the shorter lifespan of SJL/J mice relative to other common inbred strains and propensity for developing lymphomas (Chow and Ho, 1988; Yuan et al., 2009), the “aged” SJL/J mice used here are 8-12 months old. Despite the younger age of assessment, we have previously shown that the severe, non-remitting EAE phenotype of SJL/J mice is consistent when mice are ≥8 months old (Zuo et al., 2022). Detailed comparisons of SJL/J vs C57BL/6 aging metrics have not been reported, but these data suggest that some features of biological aging may be chronologically advanced in SJL/J mice. The human-to-mouse FMT portion of this study was limited to four pairs of human donors in self-reported good health, and for the scope of this study we specifically excluded individuals with any diagnosed chronic health conditions. This may explain, in part, why we observed that aged FMT donors did not universally modify EAE in mice. There is also insufficient power to draw conclusions regarding patterns in aging microbiota or metabolites, but it is perhaps logical that not all microbiotas derived from aged donors had disease-enhancing potential. Nevertheless, since our FMT pipeline introduces human-adapted microbes that elicit disparate EAE phenotypes, our approach offers a translationally relevant pipeline to mine for metabolites for treatment MS progression.

## Methods

### Animals

All experiments were performed in accordance with the ethical guidelines of the Animal Care Committees at the University of British Columbia (UBC) and University of Toronto (UofT). Conventional female or male SJL/J (SJL/JCrHsd) mice were either purchased from Envigo, USA or bred in-house at UBC or UofT (as specified in figure legends). Unless stated otherwise, all young mice used are 6-14 weeks of age, and aged mice are 8-12 months of age and age-matched across experimental groups. 12 months is the maximum age for aged mice due to high incidence of tumors in older mice (Crispens, 1973).

At UBC in the Centre for Disease Modeling, mice were housed up to 5 mice per cage in ventilated Ehret cages prepared with BetaChip bedding and had *ad libitum* access to food (PicoLab Diet 5053) and water (reverse osmosis/chlorinated (2-3 ppm)-purified). Housing rooms were maintained on a 14/10-hour light/dark cycle with temperature and humidity ranges of 20-22°C and 40-70%, respectively. Sentinel mice housed in experimental rooms were maintained on dirty bedding and nesting material and were tested on a quarterly basis for presence of mites (*Myocoptes, Radford/Myovia),* pinworm (*Aspiculuris tetaptera, Syphacia obvelata)*, fungi (*encephalitozooan cuniculi)*, bacteria (*Helicobacter spp., Clostridium piliforme, Mycoplasma pulmonis,* CAR Bacillus), and viruses (ectromelia, rotavirus, mouse hepatitis virus (MHV), murine norovirus (MNV), Minute virus of mice (MVM), lymphocytic choriomeningitis virus (LCMV), mouse adenovirus (MAV) 1/2, murine cytomegalovirus (MCMV), polyoma, pneumonia virus of mice (PVM), reovirus (REO) 3, Sendai, and Theiler’s murine encephalomyelitis virus (TMEV)).

At UofT in the Center for Cellular and Biomolecular Research, animals were housed under specific pathogen free (SPF) conditions, up to 4 mice per cage in ventilated Allentown cages prepared with bedding, nesting material, and enrichment. All mice had *ad libitum* access to 18% protein rodent diet (Teklad 2918) and acidified water. Housing rooms were maintained on a 12/12-hour light/dark cycle, temperatures of 20-26°C, and humidity of 35-70%. Sentinel animals were housed in similar conditions and tested quarterly for mites and pinworm in addition to MHV, MVM, MPV, MNV, TMEV, and rotavirus, and annually for Sendai, *M. pulmonis*, PVM, REO3, LCMV, ectromelia, and MAV1/2.

Germ-free SJL/J mice were generated through rederivation via 2-cell embryo transfer into germ-free pseudopregnant Swiss Webster females at the International Microbiome Centre (University of Calgary) and bred at UofT and UBC. GF mice were bred and housed in sterile vinyl isolators (Class Biologically Clean, Madison WI). At UBC, fecal pellets were sampled every 2 weeks to verify sterility by aerobic and anaerobic culturing, and monthly by 16S qPCR. For FMT studies, GF mice were transferred to Ehret cages, gavaged immediately with prepared samples and then housed under SPF conditions. At UofT, GF mice were bred and housed in either Tecniplast ISOcage™ cages or semi-rigid isolators. Fecal pellets were sampled every 2 weeks for monitoring germ-free status by culture or PCR for detection of bacterial 16S rRNA and fungal 18S and 28S ITS.

### Adoptive transfer experimental autoimmune encephalomyelitis

Immunization of female donor mice aged 7-9 weeks old was achieved using an emulsion containing 100 μg PLP_131-159_ and 200 μg *Mycobacterium* suspended in incomplete Freund’s adjuvant (IFA) and PBS. Donor mice were injected subcutaneously with 100 μL of emulsion at each hind flank and at the neck (UofT) or one injection in the hind flank (UBC). On day 9 following immunization, donor animals were euthanized and inguinal, axillary, brachial, and superficial cervical lymph nodes and spleens were harvested. For restimulation in culture, lymph nodes and spleens were mashed directly through a 70 μm filter into sterile ice-cold PBS. Cells were washed, counted, and resuspended in culturing media. Cells were transferred into culture flasks at 2.5 x 10^8^ cells per T-75 flask, in a final volume of 50 mL of media per flask. Cells were cultured in the flask upright for 72 hours at 37°C, 5% CO_2_ in the presence of 10 μg/mL PLP_139-151_, 10 ng/mL rIL-23, 20 μg/mL α-IL-4, and 20 μg/mL αIFNγ. Details of relevant reagents used at UBC and UofT are listed in **Table 1**.

**Table 1:** Reagents.

| Purpose/Reagent | UBC | UofT |
| --- | --- | --- |
| **EAE** |  |  |
| PLP<sub>139-151</sub> | GenScript (RP10550)<br>CanPeptide | CanPeptide |
| Mycobacterium | BD Difco | BD Difco |
| IFA | BD Difco | BD Difco |
| **Cell culture** |  |  |
| rIL-23 | R&D Systems (1887-ML-010/CF) | R&D Systems (1887-ML-010/CF) |
| αIL-4 | Bio X Cell (clone 11B11, BE0045) | Bio X Cell (clone 11B11, BE0045) |
| αIFNγ | Bio X Cell (clone XMG1.2, BE0055) | Bio X Cell (clone XMG1.2, BE0055) |
| **Antibiotics** |  |  |
| Ampicillin | Ampicillin trihydrate (Thermo, J6651422) | Ampicillin sodium salt (BioShop, AMP201) |
| Neomycin | Neomycin trisulfate salt hydrate (Sigma, N6386) | Neomycin sulfate (BioShop, NEO201) |
| Vancomycin | Vancomycin hydrochloride (Chem-Impex Intl., 00315) | Vancomycin hydrochloride (BioShop, VAN990) |
| Metronidazole | Metronidazole (Sigma, M3761) | Metronidazole (Sigma, M1547) |
| Ertapenem | Invanz® (ertapenem sodium, Merck)<br>Ertapenem sodium (generic) | Invanz® (ertapenem sodium, Merck) |
| Cefoperazone | Cefoperazone sodium salt (Sigma, C4292) | Cefoperazone sodium salt (Sigma, C4292) |
| Clindamycin | Dalacin® C (clindamycin hydrochloride capsules, Pfizer) | Antirobe® (clindamycin hydrochloride solution, Zoetis) |

Culture media: RPMI appended with 10% (v/v) FBS, 1% (v/v) GlutaMAX™, 1 mM sodium pyruvate, 1% (v/v) non-essential amino acids, 100 U/mL Penicillin-Streptomycin, and 55 µM □-mercaptoethanol.

After 72 hours, cells were collected from flasks and washed. For data shown in Fig. 7 and S2, cells were stained with CFSE (Thermo Fisher, C34554) according to manufacturer instructions. For injection into recipient mice, cells were resuspended at a final concentration of 1.0×10^8^ cells/mL. Recipient mice were either injected with 1×10^7^ or 2×10^7^ cells (indicated where relevant) intraperitoneally. Mice were evaluated for clinical health and EAE severity daily after the onset of EAE symptoms. EAE severity was scored according to a composite 16-point scale, which measures mobility impairments in each limb and the tail, as previously described(Pikor et al., 2015; Zuo et al., 2022). Each limb was graded from 0 (asymptomatic) to 3 (complete paralysis), the tail is graded from 0 (asymptomatic) to 2 (limp tail), and the righting reflex is scored from 0 to 2, with 0 assigned for normal righting reflex, 1 for slow righting reflex, and a 2 for a delay of more than 5 seconds in the righting reflex. Each criterion was measured in 0.5-point increments, thus composite scores range from 0 (completely asymptomatic) to 16 (fully quadriplegic with limp tail and significantly delayed righting reflex).

Mice were monitored for up to 25 days, and disease scores and body weight measurements were taken daily. For supportive care starting 4-5 days after adoptive transfer, cage bedding was switched to Pure-o’Cel material and cages were placed on heat pads set to 37°C. One (1)-3 subcutaneous injections of Ringer’s lactate solution were given based on degree of dehydration, and wet mash using powdered 19% protein rodent chow (Teklad, 2919) or DietGel Recovery (ClearH2O) was supplied in cages and refreshed daily. For paralyzed animals, bladders were gently expressed daily 1-3 times, as needed.

### Antibiotic treatment and FMT

For broad-spectrum antibiotic depletion, drinking water was supplemented with either 5% (w/v) sucrose for control groups or 5% sucrose with an antibiotic cocktail (ampicillin, neomycin, vancomycin, and metronidazole at 1 mg/mL). Supplemented water was given *ad libitum* starting 7 days prior to EAE induction, and continued throughout the course of EAE, refreshed every 2-3 days.

For microbiome depletion prior to microbiome transfer, SPF-housed SJL/J mice were microbiome-depleted using antibiotic cocktails as previously described (Staley et al., 2017). Briefly, antibiotics were added to animal facility drinking water, each at 1 mg/mL. A “non-absorbable” cocktail (ertapenem, neomycin, vancomycin) was administered for 7 days, followed by a 2-day wash-out with regular drinking water. Next, a “systemic” cocktail (ampicillin, cefoperazone, clindamycin) was administered for 7 days, followed by a 2-day washout with regular drinking water. A final round of “non-absorbable” cocktail was administered for 7 days, followed by a 2-day wash-out period prior to fecal/cecal microbiota transplant. Antibiotic-treated mice were orally gavaged with 100-200 μL of donor material (fecal material from human donors and cecal material from mouse donors, all herein referred to as “FMT”). Animals were transferred from standard SPF housing to CL2 housing at the time of oral gavage, which is maintained with the same environmental conditions described above. Transferred microbiome was allowed to settle for 3-5 weeks following oral gavage to allow microbiota colonization prior to any further experimental procedures. Details of specific antibiotics used at UBC and UofT are listed in **Table 1**.

### Mouse cecal material processing

Whole ceca from young or aged mice were collected in sterile 1.5 mL collection tubes and either snap-frozen on dry ice and stored at −80°C or moved immediately to an anaerobic chamber for processing. Individual ceca were transferred to a sterile homogenization tube with metal bead containing 500 µL sterile, pre-reduced PBS. Ceca were homogenized in a bead beater homogenizer for 5 minutes at 1800 rpm. Homogenate was filtered through a 70 µm nylon filter to remove debris. If several donor ceca were collected, cecal material was pooled into an age-and sex-matched batch. The cecal suspension was diluted to a total volume of 2 mL PBS per cecum. Suspensions were aliquoted into 1 mL cryovials and resuspended in 15% glycerol for storage at −80°C.

### Human fecal material processing

Stool samples were collected from donors on site at UofT (see **Table 2**). All pairs of donors were required to (1) have been born in Canada or immigrated 10+ years prior, (2) be cohabitating for the past 5 years, (3) be disparate in age by 25 years, (4) be of the same sex, and (5) be first-degree relatives. Stool was deposited into 50 mL collection tubes, and either stored at −80°C or moved into an anaerobic chamber (5% H_2_, 5% CO_2_, 90% N_2_) for immediate processing. All reagents used inside the anaerobic chamber were sterilized and degassed prior to use. An aliquot of stool was weighed and deposited into a fresh 50 mL conical tube, and sterile 0.3 mm glass beads were added. Sterile, pre-reduced PBS was added at 10 mL per 1 g of stool. Stool was gently vortexed in short bursts until homogenized, then poured through a 0.25 mm metal strainer to remove beads and large particles. Filtered homogenate was centrifuged at 5000 x *g* for 20 minutes at room temperature. Supernatant was discarded, and the pellet was weighed and resuspended at 10 mL of 15% glycerol in PBS per gram. Samples were transferred to −80°C as 1 mL aliquots for storage.

**Table 2:** Human FMT donors.

| **Donor ID** | **Sex** | **Age Difference (years)** |
| --- | --- | --- |
| Pair 1 | Male | 25 |
| Pair 2 | Female | 29 |
| Pair 3 | Male | 30 |
| Pair 4 | Female | 29 |

### Quantitative polymerase chain reaction (qPCR) for 16S rDNA

Fecal pellets and DNA extraction were performed as described above. Total DNA contained in the volume eluted from the Macherey-Nagel NucleoSoil columns were quantified by NanoDrop. 500 ng of template DNA was used for all qPCR reactions. Maxima SYBR Green/ROX qPCR Master Mix (Thermo Fisher, K0221) was combined with template DNA and primers as per manufacturer instructions. Uni926F and Uni1062R forward and reverse primers were used to detect 16S sequences (Bacchetti De Gregoris et al., 2011). A final reaction volume of 25 µL was used per reaction. A three-step cycling protocol was used as per manufacturer recommendation. Briefly, initial denaturation was performed at 95°C for 15 seconds, followed by 40 cycles of denaturation (95°C, 15 seconds), annealing (60°C, 30 seconds), and extension (72°C, 30 seconds). Mean quantification cycle (Cq) values are calculated from technical triplicates for each sample.

### 16S rRNA gene sequencing

Fresh fecal pellets from animals were collected either directly from live animals or from the distal colon at experimental endpoints. Pellets were stored at −20°C until use. Remaining human donor material was also refrozen at −20°C after thawing once for oral gavage. DNA extraction was performed using the Macherey-Nagel NucleoSoil kit according to manufacturer instructions, and DNA quality was assessed using NanoDrop. Samples that met a minimum value of 1.8 for OD_260/280_ were considered sufficiently pure for sequencing.

Sequencing was performed at the Center for Analysis of Genome Evolution and Function. The V4 hypervariable region of the 16S rRNA gene was amplified using uniquely barcoded 515F (forward) and 806R (reverse) sequencing primers (Caporaso et al., 2012). The V4 region was amplified by cycling the reaction at 95°C for 3 minutes, 18x cycles of 95°C for 15 seconds, 50°C for 15 seconds and 72°C for 15 seconds, followed by a 5-minute 72°C extension. All amplification reactions were done in duplicate to reduce amplification bias, pooled, and checked on a 1% agarose TBE gel. Pooled duplicates were quantified using PicoGreen and combined by even concentrations. The library was then purified using Ampure XP beads and loaded on to the Illumina MiSeq for sequencing, according to manufacturer instructions (Illumina, San Diego, CA). Sequencing was performed using the V2 (150bpx2) chemistry. A single-species (*Pseudomonas aeruginosa* DNA), a mock community (Zymo Microbial Community DNA Standard D6305), and a template-free negative control were included in the sequencing run.

The dada2 pipeline was used for inferring amplicon sequence variants (ASVs) from individual fastq files (Callahan et al., 2016). Briefly, the final 10 bases of all sequences were trimmed, and default filtering parameters for dada2 were used for denoising (maxN = 0, trunQ = 2, and maxEE = 2). Merging of paired reads, construction of an ASV table, and removal of chimeras were all performed through the dada2 pipeline. Taxonomy assignment was performed through the use of the Silva 138.1 NR99 reference database. dada2 outputs (ASV table, taxonomy table) and sample metadata were combined and analyzed further using the phyloseq package. ASVs not assigned to a known phylum and low prevalence taxa were removed prior to all downstream analyses. R packages used are listed in **Table 3**.

**Table 3:** R packages used for data visualization and statistical testing.

| Package | Citation | Data | Purpose |
| --- | --- | --- | --- |
| dada2 | (Callahan et al., 2016) | 16S | Pre-processing of raw sequencing data |
| phyloseq | (McMurdie and Holmes, 2013) | 16S | Handling and analysis of microbiome data |
| ampvis2 | (Andersen et al., 2018) | 16S | Heatmap visualizations |
| vegan | (Oksanen et al., 2025) | 16S | Statistical comparisons (PERMANOVA, PERMDISP2) |
| pairwiseAdonis | (Martinez Arbizu, 2020) | 16S | Statistical comparisons (pairwise PERMANOVA) |
| tidyverse | (Wickham et al., 2019) | 16S, metabolites | Data visualization and plot generation |
| DESeq2 | (Love et al., 2014) | 16S | Differential abundance testing |

### Sample preparation for metabolomic analysis

For metabolite extraction, whole, snap-frozen mouse serum, cecal content, fecal pellet, cortex, and leptomeninges samples were processed as follows. Frozen cortex samples were crushed and ground into a fine powder with a pre-chilled mortar and pestle, with dry ice added throughout to prevent sample thawing. Then, 10 μL methanol was added per mg of powdered sample, and samples were sonicated for 5 min. Extracts were pelleted at 5,000 x *g* for 5 min, and supernatants were transferred to another 20 mL glass vial. Remaining pellets were further extracted with another 10 min of vigorous stirring in 5 mL ethanol. The supernatants were combined and then dried in a SpeedVac (ThermoFisher Scientific) vacuum concentrator. Dried materials were resuspended in 300 μL of methanol. Samples were pelleted at 5,000 g for 5 min and clarified extracts were transferred to fresh HPLC vials and stored at −20°C until analysis.

For the analysis of both donor and mice serum samples, 400 μL of methanol was added per 100 μL of sample. The samples were vortexed and centrifuged, and supernatant was collected. An additional volume of 200 μL of methanol was added to the residue and the vial was again vortexed and centrifuged. The combined supernatants were dried in a SpeedVac (ThermoFisher Scientific) vacuum concentrator. Samples were resuspended in 50 μL of methanol and pelleted at 5,000 g for 5 min. Clarified extracts were transferred to fresh HPLC vials and stored at −20°C until analysis.

### Mass spectrometric analysis

HPLC-HRMS analysis was conducted using a Thermo Fisher Scientific Vanquish Horizon ultra-HPLC (UHPLC) System coupled to a Thermo Q Exactive hybrid quadrupole-orbitrap high-resolution mass spectrometer equipped with a heated electrospray ionization (ESI) ion source, using Thermo Scientific Xcalibur software (v.4.3.73.11). For each sample, 1 μL of the above prepared methanol extracts was injected and separated using a water-acetonitrile gradient on a Thermo Scientific Accucore C18 column (150 × 2.1 mm, 1.5 μm particle size) and maintained at 40°C. Optima grade solvents were purchased from Fisher Scientific. Solvent A was 0.1% formic acid in water and solvent B was 0.1% formic acid in acetonitrile. A/B gradient started at 1% B for 3 min, then increased linearly to 98% B at 20 min, followed by 98% B for 5 min, then down to 1% B for 3 min. Mass spectrometer parameters: +3.5 kV spray voltage, 380°C capillary temperature, 40°C probe heater temperature, 60 AU sheath flow rate, 20 AU auxiliary flow rate, 2 AU sweep gas; S-lens radio frequency level 50.0, resolution 60,000 at m/z 200, m/z range 80–1,200, AGC target 3E6. Normalized collision energy was set to a stepwise increase from 10 to 30 to 50 % with z = 1 as default charge state. For MS2 data analysis, raw spectra were converted to .mzXML files using MSconvert (ProteoWizard). MS1 and MS2 feature extraction was performed using MZmine 3.4.14 (Schmid, R et al. 2023). The output was further processed through GNPS for molecular networking and library matching. Chemical feature prioritization and identification of chemical drivers was performed using Metaboseek (Helf, M.J et al. 2022), applying filters on fold change with threshold 3 and p-value with threshold 0.05. The acquired data and workflows are available under the FAIR (Findability, Accessibility, Interoperability, and Reuse) principles. Specifically, MS/MS data are stored in the MassIVE repository (<u>massive.ucsd.edu</u>) under identifier MSV000096116.

### In vitro and in vivo IPA treatment

Splenocytes were prepared for culture as described above. In a 96-well U-bottom plate, 1.0×10^6^ cells were plated per well. IPA (Sigma 57400) solubilized in DMSO was added to a final concentration of 200 μM. The same volume of DMSO was added to control wells. After 72 hours, cells were analyzed by flow cytometry.

For in vivo treatment, mice were given daily systemic injections of 20 mg/kg IPA suspended in sterile PBS or vehicle control as previously described (Rothhammer et al., 2016). Injections alternated between i.p. and s.c. (UBC) or were daily i.p. (UofT). Treatment began 7 days prior to A/T EAE and continued throughout the course of the experiment. Suspensions were generated fresh daily immediately before injection.

### Sample preparation for histology analyses

At experimental endpoints, mice were euthanized by CO_2_ overdose and perfused intracardially with ice-cold PBS. Whole brains were dissected out, cut sagittally, and one hemisphere was directly deposited into 10% formalin. Following at least overnight fixation, samples are sent to The Center for Phenogenomics for embedding into paraffin blocks and sectioning into 8 µm sections.

For Hematoxylin and Eosin staining, sections were first dewaxed in xylene and rehydrated in sequentially lower concentrations of ethanol. Slides were submerged in Hematoxylin (Sigma, MHS32) solution for 10 minutes, then rinsed under slowly running tap water briefly. Sections were then immersed in 100% ethanol for 5 minutes, then stained with Eosin Y (Sigma E4382) solution for 3 minutes. Slides were dipped in 90% ethanol to differentiate Eosin staining, then dehydrated in sequentially increasing concentrations of ethanol and xylene. Cover slips were mounted with Entellan and allowed to dry before imaging.

For immunocytochemistry, sections were dewaxed and rehydrated as above. Slides were immersed in 0.3% hydrogen peroxide in methanol for 20 minutes to quench endogenous peroxidase activity. Antigen retrieval was performed using Tris-EDTA buffer (10 mM Tris base, 1 mM EDTA, adjusted to pH of 9.0). Slides were blocked using Fc block (BioXCell BE0307-25MG-A) in 10% normal goat serum and 5% bovine serum albumin (BSA) in 0.1% PBS-Tween 20. Iba1 antibody (abcam ab178846) was diluted 1:4000 from stock in 5% BSA in 0.1% PBS-Tween 20. Primary antibody was incubated overnight at 4°C in a humidity chamber. The BrightVision, 2 steps detection system (Immunologic, DPVB500HRP) was used for detection of primary antibody. The Signalstain® DAB Substrate Kit (Cell Signaling Technology, 8059S) was used for development of signal.

For analysis, all stained sections were scanned using the Zeiss Axioscan 7. All images were acquired at 20X magnification. Quantifications were performed using the open source software QuPath (Bankhead et al., 2017).

### Sample preparation for flow cytometry

For brain and spinal cord, tissues were mechanically homogenized through a 70 µm mesh filter into ice-cold CMF-HBSS appended with 10 mM HEPES, 150 mM NaCl, 1 mM MgCl_2_, 5 mM KCl, and 1.8 mM CaCl_2_. DNase I and collagenase D were added to final concentrations of 60 ug/mL and 1 mg/mL, respectively. Tissue was digested at 37°C on an orbital shaker for 30 minutes. EDTA was added at 1 mM to halt enzyme activity. Myelin was separated from the cell suspension using 30% isotonic Percoll, centrifugating at 800 x *g* for 20 minutes without brakes. Cells were washed in PBS, followed by antibody staining.

For spleens and lymph nodes, tissues were mechanically homogenized through a 70 µm mesh filter into ice-cold PBS. ACK buffer was added to spleen samples only, to lyse red blood cells. Cells were washed in PBS, followed by antibody staining.

Cell viability staining (ThermoFisher, L34965 or L34955) was done in PBS before antibody staining. All antibody staining was done in PBS with 2% (v/v) FBS. For intracellular targets, cells were fixed, permeabilized, and stained using the Foxp3/Transcription Factor Staining Buffer Set (Invitrogen, 00-5523-00) following manufacturer instructions

### Statistics

Data visualizations and statistical analysis were performed either using GraphPad Prism, RStudio, QuPath, or FlowJo software. Relevant statistical tests are specified in figure captions. Relevant R packages are described in **Table 3**.

## Supplemental materials

Details of all differential abundance testing or statistical testing for 16S rDNA sequencing are presented in Table S1. Metabolites identified and shown in Fig. 6 are listed in Table S2. The parameters used in MZmine3 and GNPS (Wang et al., 2016) are listed in Table S3.

## Resource Availability

This study did not generate new unique reagents. Raw 16S rDNA sequencing files can be found in the NCBI Sequence Read Archive under BioProject accession number PRJNA1370512. Metabolomic data and workflows are available under the FAIR (Findability, Accessibility, Interoperability, and Reuse) principles. MS/MS data are stored in the MassIVE repository (<u>massive.ucsd.edu</u>) under identifier MSV000096116. The molecular network output and its parametrization can be accessed on the GNPS repository at: https://tinyurl.com/Pu-Fettig-metabolite-GNPS. Any additional information required to reanalyze the data reported in this paper is available from the lead contact upon request.

## Acknowledgements

The authors are grateful for support from MS Canada (AWD-009454 MSSCA 2019 & AWD-023455 MSSCA 2022 to JLG & LCO), the Canadian Institutes for Health Research (FDN-159922 to JLG), Praespero: An Autoimmune Research Fund (JLG), the Research Corporation for Science Advancement Scialog award (RCSA-28637 to LCO), Michael Smith Health Research BC (RT-2023-3183 to LSH), and the MS Canada endMS Personnel Award program (AP, NMF, IN, MZ, SJP). JLG and LCO received salary support through the Canada Research Chairs program. We thank veterinary staff and animal care technicians at the International Microbiome Center (University of Calgary), DCM (University of Toronto), and the Centre for Disease Modelling (UBC); cytometry experts at the Faculty of Medicine Flow Cytometry core facility (University of Toronto) and the Life Sciences Institute ubcFLOW core (UBC); and members of the International MS Microbiome Study for invaluable discussions and feedback. UBC core facilities are supported by the UBC GREx Biological Resilience Initiative. Work in the Gommerman and Osborne labs takes place on the traditional, ancestral, and unceded territories of the Huron-Wendat, Seneca, Mississaugas of the Credit and xwməqkwəýəm (Musqueam) First Nations.

---

**Figure S1.**
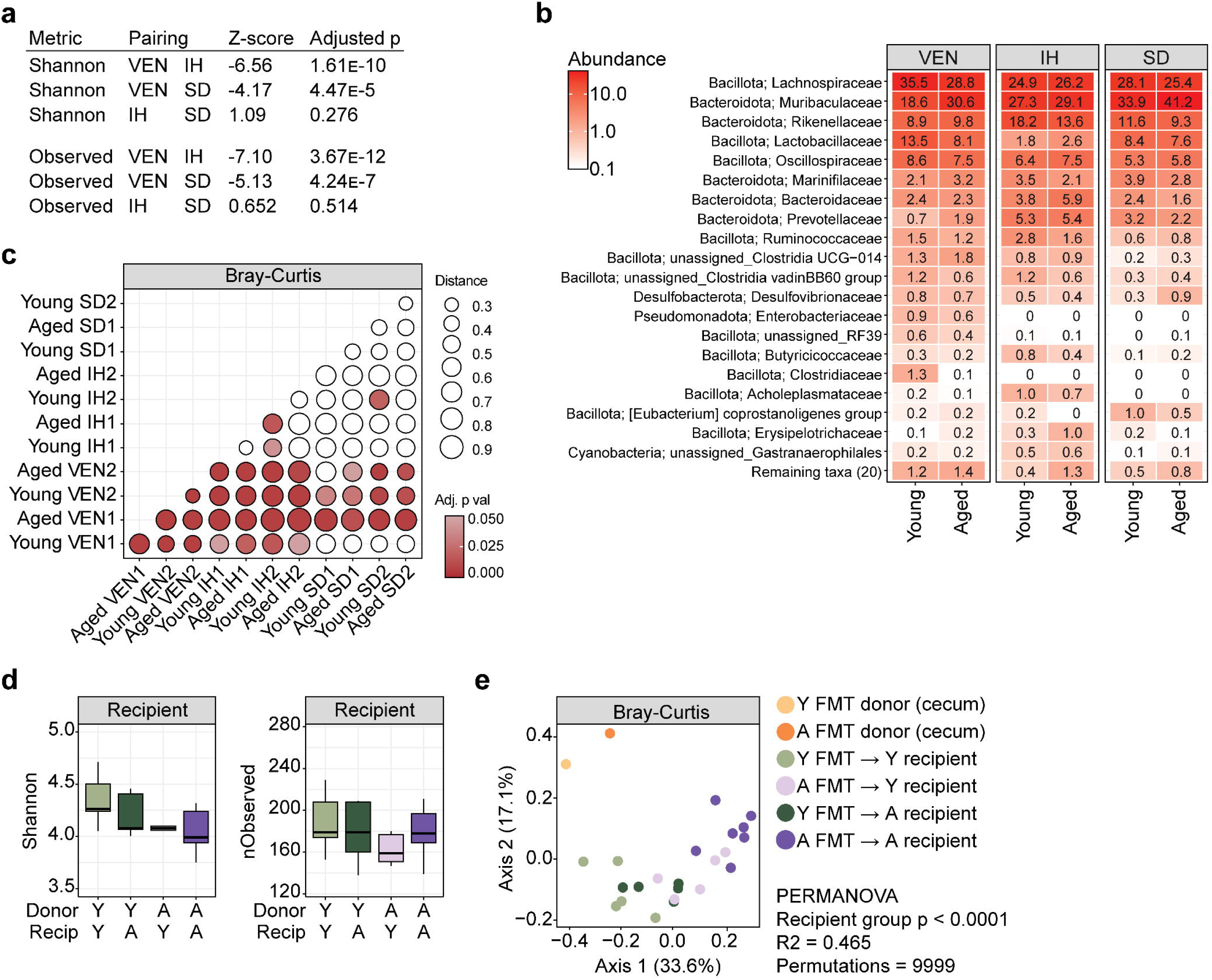
Age does not drive significant microbiome modulation in SJL/J mice. (a) Statistical comparison of cohorts with combined young and aged mice. Kruskal-Wallis test. (b) Relative abundance of top 20 most abundant families and combined remaining families. Colours indicate log_10_ abundance value. (c) Comparison of Bray-Curtis distances between young and aged mice across cohorts. Bubble size indicates mean Bray-Curtis distance, colour indicates adjusted p value. Pairwise PERMANOVA with Benjamini-Hochberg p value adjustment. (d) Shannon diversity and number of observed ASVs from CMT recipient mouse fecal material. Kruskal-Wallis test with Dunn multiple comparisons test (recipient fecal samples only). (e) PCoA plot of Bray-Curtis distances.

**Figure S2.**
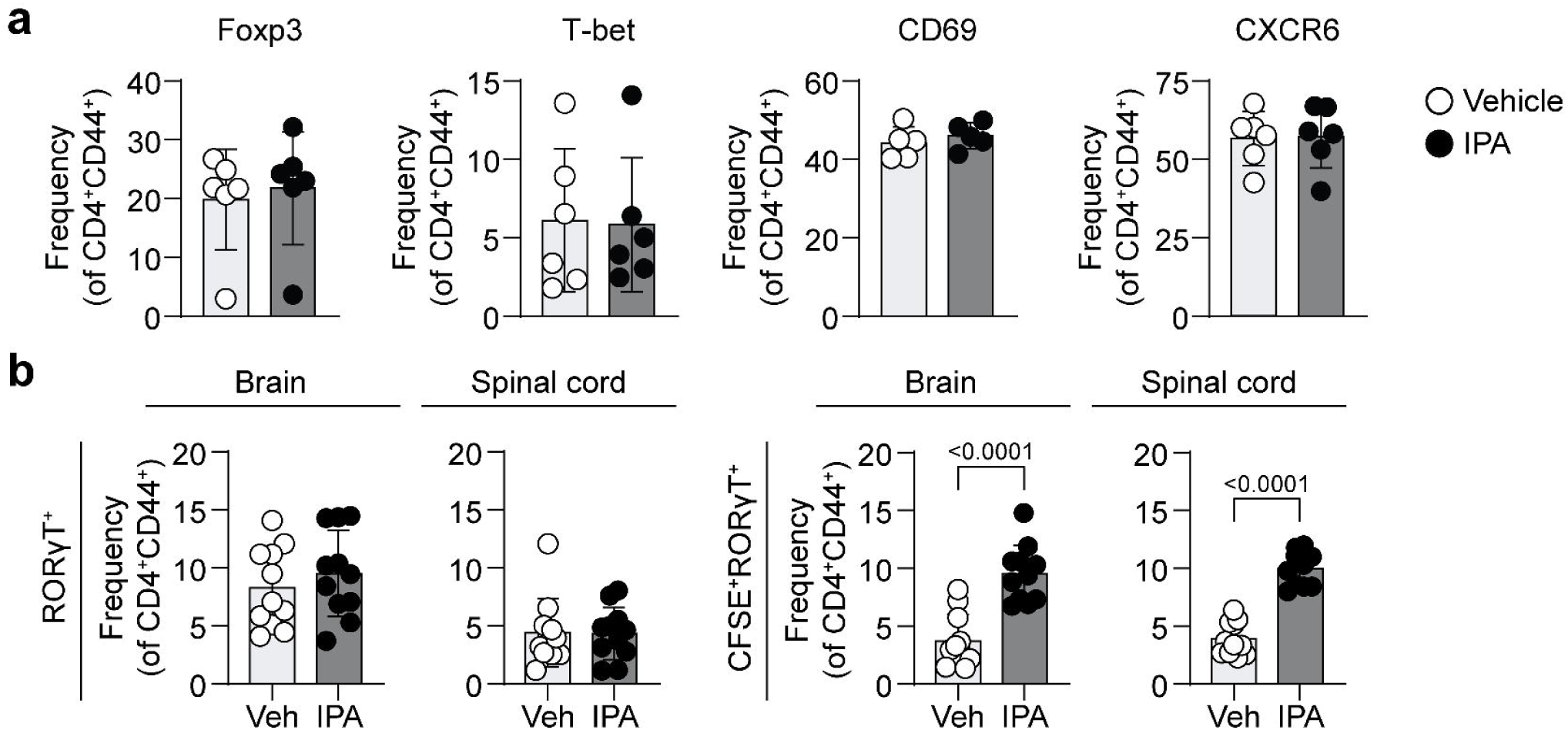
IPA drives resolution of clinical EAE and inhibits T cell IL-17A production. (c) Frequency of CNS-penetrating CD4^+^CD44^+^ T cells expressing Foxp3, T-bet, CD69, and CXCR6. (b) Frequency of total or CFSE^+^ CD4^+^CD44^+^RORgT^+^ T cells in brain and spinal cord of mice receiving IPA or vehicle-treated donor cells at day 7 post-A/T. Wilcoxon rank sum test for (a), (b).

