## Supplemental Table 3 (S3) for "Mining for disease-associated microbial metabolites in an age-dependent model of multiple sclerosis"

Supplementary Table S3. Parameters used for the MZmine and GNPS analyses.

| MZmine | Raw data  method> Feature detection | >mass detection | Set filters | | |  | | MS1 | | | | MS2 |
| --- | --- | --- | --- | --- | --- | --- | --- | --- | --- | --- | --- | --- |
|  |  |  |  |  |  | Scan number | |  | | | |  |
|  |  |  |  |  |  | Base Filtering Integer | |  | | | |  |
|  |  |  |  |  |  | Retention time | |  | | | |  |
|  |  |  |  |  |  | MS level | | 1 | | | | 2 |
|  |  |  |  |  |  | Scan definition | |  | | | |  |
|  |  |  |  |  |  | Polarity | | ANY | | | | ANY |
|  |  |  |  |  |  | Spectrum type | | ANY | | | | ANY |
|  |  |  | Mass detector | | | Mass detector | | CENTROID | | | | CENTROID |
|  |  |  |  |  |  | Noise level | | 100 | | | | 50 |
|  |  | >ADAP  Chromatogram Builder | Set filter | | | Scan number | |  | | | | |
|  |  |  |  |  |  | Base Filtering Integer | |  | | | | |
|  |  |  |  |  |  | Retention time | |  | | | | |
|  |  |  |  |  |  | MS level | | 1 | | | | |
|  |  |  |  |  |  | Scan definition | |  | | | | |
|  |  |  |  |  |  | Polarity | |  | | | | |
|  |  |  |  |  |  | Spectrum type | |  | | | | |
|  |  |  | Min group size in # of scans | | | | | 5 | | | | |
|  |  |  | Group intensity threshold | | | | | 100000 | | | | |
|  |  |  | Min highest intensity | | | | | 300000 | | | | |
|  |  |  | m/z tolerance | | | | | 0.02 m/z or 10 ppm | | | | |
|  | >Feature list methods | >Feature detection> Chromatographic deconvolution | Algorithm> Local minimum search | | Chromatographic threshold | | | | | | | 87 |
|  |  |  |  |  | Search minimum in RT range (min) | | | | | | | 0.20 |
|  |  |  |  |  | Minimum relative height | | | | | | | 0.0 |
|  |  |  |  |  | Minimum absolute height | | | | | | | 300000 |
|  |  |  |  |  | Min ratio of peak top/edge | | | | | | | 2 |
|  |  |  |  |  | Peak duration range (min) | | | | | | 0.01 – 5.00 | |
|  |  |  | m/z range for MS2 scan pairing (Da) | | | | | | ☑ 0.025 | | | |
|  |  |  | RT range for mS2 scan pairing (min) | | | | | | ☑ 0.15 | | | |
|  |  | >Isotopes> Isotopic peak  grouper | m/z tolerance | | | | | | 0.015 m/z or 3 ppm | | | |
|  |  |  | Retention time tolerance | | | | | | 0.03 | | | |
|  |  |  | Maximum charge | | | | | | 2 | | | |
|  |  |  | Representative isotope | | | | | | Most intense | | | |
|  |  | >Alignment>  Join Aligner | m/z tolerance | | | | | | 0.015 m/z or 5 ppm | | | |
|  |  |  | Weight for m/z | | | | | | 3 | | | |
|  |  |  | Retention time tolerance | | | | | | 0.20 | | | |
|  |  | >Filtering> Feature list row filter | | Reset the peak number ID | | | | | ☑ | | | |
|  |  | Peak finder (multithreated) | | | | | m/z tolerance | | | 0.002 m/z , 10 ppm | | |
| GNPS | Feature networking | Basic options | Precursor Ion Mass Tolerance | | | | | | 0.02 | | | |
|  |  |  | Fragment Ion Mass Tolerance | | | | | | 0.02 | | | |
|  |  | Advanced Network Options | Min Pairs Cos | | | | | | 0.7 | | | |
|  |  |  | Minimum Matched Fragments Ions | | | | | | 6 | | | |
|  |  |  | Maximum shift between precursors | | | | | | 500 | | | |
|  |  |  | Network TopK | | | | | | 10 | | | |
|  |  |  | Maximum Connected Component Size | | | | | | 100 | | | |
